# Nipponbare and wild rice species as unexpected tolerance and susceptibility sources against *Schizotetranychus oryzae* (Acari: Tetranychidae) mite infestation

**DOI:** 10.1101/2020.01.22.914184

**Authors:** Giseli Bufon, Édina Aparecida dos Reis Blasi, Thainá Inês Lamb, Janete Mariza Adamski, Joséli Schwambach, Felipe Klein Ricachenevsky, Amanda Bertolazi, Vanildo Silveira, Mara Cristina Barbosa Lopes, Raul Antonio Sperotto

## Abstract

Cultivated rice (*Oryza sativa* L.) is frequently exposed to multiple stresses, including *Schizotetranychus oryzae* mite infestation. Rice domestication has narrowed the genetic diversity of the species, reducing the stress resistance and leading to a wide susceptibility. Therefore, wild rice species present an alternative to search for this lost variability. Aiming to observe the response of two wild rice species (*Oryza barthii* and *Oryza glaberrima*) and two *Oryza sativa* genotypes (cv. Nipponbare and *O. sativa* f. *spontanea*) to *S. oryzae* infestation, we used agronomic, physiological and molecular analyses. Surprisingly, analyses of leaf damage, histochemistry, chlorophyll concentration and chlorophyll fluorescence showed that the wild species present higher level of leaf damage, increased accumulation of H_2_O_2_ and lower photosynthetic capacity when compared to *O. sativa* genotypes under infested conditions. Infestation did not affect plant height, but decreased tiller number, except in cv. Nipponbare, whose development was not affected. Infestation also caused the death of wild plants during the reproductive stage, unlike *O. sativa* genotypes, which were able to tolerate stress and produce seeds. While infestation did not affect the weight of 1,000 grains in both *O. sativa* genotypes, the number of panicles per plant was affected only in *O. sativa* f. *spontanea*, and the percentage of full seeds per panicle and seed length were increased only in cv. Nipponbare. Proteomic analysis allowed us to identify 195 differentially abundant proteins when comparing susceptible (*O. barthii*) and tolerant (*O. sativa* cv. Nipponbare) genotypes under control and infested conditions. We found that *O. barthii* has a less abundant antioxidant arsenal. In addition, it is unable to modulate proteins involved with general metabolism and energy production under infested condition. In Nipponbare we found high abundance of detoxification-related proteins, general metabolic processes and energy production, which allows us to suggest that, under infested condition, the primary metabolism is maintained more active compared to *O. barthii*. Also, Nipponbare presents a greater abundance of defense-related proteins, such as osmotin, ricin B-like lectin, and protease inhibitors of the Bowman Birk trypsin inhibitor family, as well as higher levels of the compatible osmolyte Proline under infested condition. Identification of these differentially abundant proteins can be used as an important biotechnological tool in breeding programs that aim increased tolerance to phytophagous mite infestation.

## Introduction

Rice (*Oryza sativa* L.) is extremely important for human nutrition, representing around 20% of the daily intake calories in the world and considered a staple food for over half of the world’s population (Muthayya et al. 2014). To meet this nutritional demand, natural resources must be used efficiently and crop production needs to be increased (Godfray et al. 2010). However, the search for new agricultural frontiers and the availability of natural resources is limited, exposing rice cultivation to several disturbances caused by biotic and abiotic stresses which negatively interfere with grain yield (Palmgren et al. 2015). Among these important stresses that influence plant yield, the presence of arthropods (Oerke et al. 2006) impact about 18% to 26% of annual crop production worldwide, causing losses of over US$ 470 billion (Culliney et al. 2014). Most of these losses (13-16%) occur in the field before harvesting, and losses are most often described in developing countries (Culliney et al. 2014). One of the arthropods that attack rice crops in Brazil, and has been reported in several South American countries, is *Schizotetranychus oryzae* Rossi de Simons mite species, which can cause more than 60% loss in rice grain yield (Buffon et al. 2018). An interesting strategy for reducing these losses is the search for mite-tolerant cultivars, as these plants can tolerate mite infestation without decreasing their grain yield and without the need for acaricides (Sperotto et al. 2018a).

The search for genetic variability in the *Oryza* genus is a shortcut to obtain resistant/tolerant rice genotypes and to develop cultivars that adapt to natural adversities (Menguer et al. 2017; Szareski et al. 2018). Several authors report wild rice plants that are resistant/tolerant to biotic and abiotic factors (Brar and Khush, 2003). The use of wild rice species in breeding programs can facilitate adaptation to biotic and abiotic stresses, as well as meeting the demand for food security in the current scenario of rapidly growing world population (Henry, 2014). The *Oryza* genus is composed of 24 species, two cultivated (*O. sativa* and *O. glaberrima*) and other 22 wild species. These wild species represent between 15 and 25 million years of evolutionary diversification (Vaughan, 1994; Menguer et al. 2017).

*O. barthii* is an annual wild African rice that is commonly known as the progenitor of *O. glaberrima*, the rice cultivated species grown in Africa. *O. barthii* is tolerant to several biotic and abiotic stresses (Khush, 1997), being adapted to more adverse ecological conditions, and resistant to multiple environmental restrictions (Sarla and Swamy, 2005). Another genotype of the *Oryza* genus that show tolerance to environmental stresses is red rice (*Oryza sativa* f. *spontanea*), which is the result of rice de-domestication process. Therefore, the genetic background, morphology and growth behavior is similar to cultivated rice (Qiu et al. 2017). However, differences in tolerance to stressful conditions may occur during de-domestication, e. g. red rice has tolerance to cold, high salinity and drought, along with blast resistance and positive germination characteristics (Dai et al. 2014). Due to its greater competitiveness and stress tolerance, red rice has become one of the most feared and harmful weeds in rice producing regions worldwide (Di et al. 2011). Nonetheless, interesting traits found in red rice can be eventually transferred to cultivated genotypes.

Aiming to identify novel resistance/tolerance mechanisms in rice plants exposed to *S. oryzae* mite infestation, we analyzed two wild species (*O. barthii* and *O. glaberrima*), red rice (*O. sativa* f. *spontanea*) and cv. Nipponbare (*O. sativa* ssp. *japonica*). Although we expected wild rice species to show higher tolerance to mite infestation, we found unexpected results. According to our data, both wild rice species are extremely sensitive to mite infestation, while cv. Nipponbare seems to be more tolerant in comparison. Therefore, we tried to understand the molecular and physiological mechanisms behind Nipponbare tolerance and wild species susceptibility to this mite. Our results may be useful for future breeding programs aiming at resistance/tolerance to phytophagous mite *S. oryzae* infestation.

## Material and methods

### Plant growth conditions and mite infestation

Seeds of *O. sativa* cv. Nipponbare and *O. sativa* f. *spontanea* (hereafter “Nipponbare” and “red rice”, respectively), *O. glaberrima* and *O. barthii* species were surface sterilized and germinated for four days in an incubator (28°C) on paper soaked with distilled water. After germination, plantlets were transferred to vermiculite/soil mixture (1:3) for additional 14 days in greenhouse conditions, and then transferred to plastic buckets containing soil and water. Plastic buckets containing rice plants highly infested by *S. oryzae* were kindly provided by Instituto Rio-Grandense do Arroz (IRGA, Cachoeirinha, RS), and were used to infest rice plants in our experiment. Fifty plants (V10-13 stage, according to Counce et al., 2000) of each cultivar/species (five plants per bucket) were infested by proximity with the bucket containing the highly infested plants placed in the center of the other buckets. For greater homogeneity of infestation and contact, buckets of each cultivar were rotated at a 90° angle counterclockwise every two days. Ten plants of each cultivar were cultivated without infestation (control condition).

The level of damage caused by *S. oryzae* was analyzed from V10-13 stage until the plants reach its final stage of reproductive development (panicle maturity, R9 stage; Counce et al. 2000). Evaluation of damage in the abaxial and adaxial faces of leaves was based on a classification of four levels of infestation: control condition, without any sign of infestation; early infested (EI) leaves, 10 to 20% of damaged leaf area, average of 168 hs of exposure to the mite; intermediate infested (II) leaves, 40 to 50% of damaged leaf area, average of 360 hs; and late infested (LI) leaves, more than 80% of damaged leaf area, average of 720 hs, according to Supplementary Figure 1.

### Plant height and tiller number

Plant height and tiller number were evaluated (n = 50) during the vegetative stage (V10-13, before being infested) and during the last reproductive stage (R9, control and infested plants).

### Chlorophyll a fluorescence transients

The chlorophyll a fluorescence transient was measured (n = 10) on the third upper leaves of control and infested plants in two different exposure times, one week (which correspond to EI), and 8 weeks (which correspond to LI), using a portable fluorometer (OS30p, Optisciences, UK). Before the measurements, plants were dark adapted for 20 minutes and the fluorescence intensity was measured by applying a saturating pulse of 3,000 µmol photons m^-2^ s^-1^ and the resulting fluorescence of the chlorophyll a measured from 0 to 1 s. The chlorophyll fluorescence intensity rises from a minimum level (the O level, F_O_ = 160 μs) to a maximum level (the P level, F_P_ = 300 ms), via two intermediate steps labeled J (F_J_ = 2 ms) and I (F_I_ = 30 ms) phases (Stirbet and Govindjee, 2011), also known as OJIP curve (Strasser et al. 2000).

### Total chlorophyll concentration

Samples (n = 3) containing 100 mg of leaves from rice plants submitted to control or LI were ground in liquid nitrogen and chlorophyll extracted in 85% (v/v) acetone. Chlorophyll *a* and *b* were quantified by measuring absorbance at 663 and 645 nm, and the concentrations calculated according to Ross (1974).

### In situ histochemical localization of H_2_O_2_

*In situ* accumulation of H_2_O_2_ in control and LI leaves was detected by histochemical staining with diaminobenzidine (DAB), according to Shi et al. (2010), with minor modifications. For H_2_O_2_ detection, leaves were excised and immersed in DAB solution (1 mg ml^-1^, pH 3.8) in 10 mM phosphate buffer (pH 7.8), and incubated at room temperature for 8 h in the light until brown spots were visible, which are derived from the reaction of DAB with H_2_O_2_. Leaves were bleached in boiling concentrated ethanol to visualize the brown spots, which were kept in 70% ethanol for taking pictures with a digital camera coupled to a stereomicroscope.

### Seed analysis

Seeds from Nipponbare and red rice were collected in R9 stage and the following agronomical parameters were evaluated: number of seeds (empty + full) per panicle, number of panicles per plant, percentage of full seeds per plant, weight of 1,000 full seeds, and seed length. Yield reduction caused by *S. oryzae* infestation was calculated using the following equation for each cultivar and each condition (control and infested): number of seeds (empty + full) per plant X percentage of full seeds X weight of one seed = seed weight per plant.

### Proline quantification

Control and LI leaves of Nipponbare and red rice were collected and proline accumulation was quantified according to Shukla et al. (2012). 250 mg of sample was sprayed in liquid nitrogen, homogenized in 2 ml of sulfosalicylic acid (3%) and centrifuged at 10,000 g for 30 min. Then, 1 ml of the supernatant was transferred to a new tube and 1 ml of glacial acetic acid and 1 ml of ninhydrin (1.25 g of ninhydrin + 30 ml of glacial acetic acid + 20 ml of 6 M phosphoric acid) were added. The homogenized mixture was boiled in a water bath at 100°C for 30 minutes. Subsequently, the reaction was cooled on ice and 2 ml of toluene was added, mixing in the vortex for 30 seconds. The upper phase containing proline was measured on the spectrophotometer at 520 nm. The proline level (mmol g^-1^ fresh weight) was quantified using L-proline as standard. The standard curve was prepared from a stock solution of proline (5 mM) in sulfosalicylic acid for a concentration range between 10 and 320 mmol ml^-1^. The free proline concentration of each sample was calculated from the linear regression equation obtained from the standard curve. The procedure for constructing the standard curve was the same as for free proline determination of the samples.

### Plant protein extraction and quantification

Three biological samples (250 mg of fresh matter) of control and EI leaves from *O. barthii* (mite-sensitive) and Nipponbare (mite-tolerant), each containing three leaves from three different plants, were subjected to protein extraction using Plant Total Protein Extraction Kit (Sigma-Aldrich). The protein concentration was measured using 2-D Quant Kit (GE Healthcare, Piscataway, NJ, USA).

### Protein digestion

For protein digestion, three biological replicates of 100 µg of proteins from *O. barthii* and Nipponbare leaves were used. Before the trypsin digestion step, protein samples were precipitated using the methanol/chloroform methodology to remove any detergent from samples (Nanjo et al. 2012). Then, samples were resuspended in Urea 7 M and Thiourea 2 M buffer and desalted on Amicon Ultra-0.5 3 kDa centrifugal filters (Merck Millipore, Germany). Filters were filled to maximum capacity with buffers and centrifuged at 15,000 g for 10 min at 20°C. The washes were performed twice with Urea 8 M and then twice with 50 mM ammonium bicarbonate (Sigma-Aldrich) pH 8.5, remaining approximately 50 μL per sample after the last wash. The methodology used for protein digestion was as previously described (Calderan-Rodrigues et al. 2014). For each sample, 25 μL of 0.2% (v/v) RapiGest® (Waters, Milford, CT, USA) was added, and samples were briefly vortexed and incubated in an Eppendorf Thermomixer® at 80°C for 15 min. Then, 2.5 μL of 100 mM DTT (GE Healthcare) was added, and the tubes were vortexed and incubated at 60°C for 30 min under agitation. Next, 2.5 μL of 300 mM iodoacetamide (GE Healthcare) was added, and the samples were vortexed and then incubated in the dark for 30 min at room temperature. The digestion was performed by adding 20 μL of trypsin solution (50 ng/μL; V5111, Promega, Madison, WI, USA) prepared in 50 mM ammonium bicarbonate, and samples were incubated at 37°C during 15 hs. For RapiGest® precipitation and trypsin activity inhibition, 10 μL of 5% (v/v) trifluoroacetic acid (TFA, Sigma-Aldrich) was added and incubated at 37°C for 30 min, followed by a centrifugation step of 20 min at 16,000 g. Samples were transferred to Total Recovery Vials (Waters).

### Mass spectrometry analysis

A nanoAcquity UPLC connected to a Synapt G2-Si HDMS mass spectrometer (Waters, Manchester, UK) was used for ESI-LC-MS/MS analysis. Fist was performed a chromatography step by injecting 1 μl of digested samples (500 ng/μl) for normalization to relative quantification of proteins. To ensure standardized molar values for all conditions, normalization among samples was based on stoichiometric measurements of total ion counts of MS^E^ scouting runs prior to analyses using the ProteinLynx Global SERVER v. 3.0 program (PLGS; Waters). Runs consisted of three biological replicates. During separation, samples were loaded onto the nanoAcquity UPLC 5μm C18 trap column (180 μm × 20 mm) at 5 μl/min during 3 min and then onto the nanoAcquity HSS T3 1.8 μm analytical reversed phase column (75 μm × 150 mm) at 400 nL/min, with a column temperature of 45°C. For peptide elution, a binary gradient was used, with mobile phase A consisting of water (Tedia, Fairfield, Ohio, USA) and 0.1% formic acid (Sigma-Aldrich), and mobile phase B consisting of acetonitrile (Sigma-Aldrich) and 0.1% formic acid. Gradient elution started at 7% B, then ramped from 7% B to 40% B up to 91.12 min, and from 40% B to 99.9% B until 92.72 min, being maintained at 99.9% until 106.00 min, then decreasing to 7% B until 106.1 min and kept 7% B until the end of experiment at 120.00 min. Mass spectrometry was performed in positive and resolution mode (V mode), 35,000 FWHM, with ion mobility, and in data-independent acquisition (DIA) mode; ion mobility separation (HDMS^E^) using IMS wave velocity of 600 m/s, and helium and IMS gas flow of 180 and 90 ml/min respectively; the transfer collision energy ramped from 19 V to 55 V in high-energy mode; cone and capillary voltages of 30 V and 2750 V, respectively; and a source temperature of 70°C. In TOF parameters, the scan time was set to 0.5 s in continuum mode with a mass range of 50 to 2,000 Da. The human [Glu1]-fibrinopeptide B (Sigma-Aldrich) at 100 fmol/μl was used as an external calibrant and lock mass acquisition was performed every 30 s. Mass spectra acquisition was performed by MassLynx v4.0 software.

### Bioinformatics analysis

Spectra processing and database searching conditions were performed by Progenesis QI for Proteomics Software V.2.0 (Nonlinear Dynamics, Newcastle, UK). The analysis used the following parameters: Apex3D of 150 counts for low energy threshold, 50 counts for elevated energy threshold, and 750 counts for intensity threshold; one missed cleavage, minimum fragment ion per peptide equal to two, minimum fragment ion per protein equal to five, minimum peptide per protein equal to two, fixed modifications of carbamidomethyl (C) and variable modifications of oxidation (M) and phosphoryl (STY), and a default false discovery rate (FDR) value at a 1% maximum, peptide score greater than four, and maximum mass errors of 10 ppm. The analysis used the *Oryza sativa* protein databank from Phytozome (https://phytozome.jgi.doe.gov/). Label-free relative quantitative analyses were performed based on the ratio of protein ion counts among contrasting samples. After data processing and to ensure the quality of results, only proteins present or absent (for unique proteins) in three out of three runs were accepted and submitted to differentially abundance analysis. Proteins were considered up-regulated if the fold change (FC) was greater than 1.5 and down-regulated if the FC was less than 0.6667, and both with significantly P-value ANOVA (P < 0.05).

### Statistical analysis

Data were analyzed using the Student’s *t* test (P ≤ 0.05 and 0.01) or One-Way ANOVA followed by Tukey test (P ≤ 0.05), using SPSS Base 23.0 for Windows (SPSS Inc., USA).

## Results

### Different responses of rice cultivars/species to S. oryzae infestation

Unexpectedly, we verified that wild rice species (*O. barthii* and *O. glaberrima*) and red rice (*O. sativa* f. *spontanea*) presented high susceptibility to *S. oryzae* infestation, while *O. sativa* cv. Nipponbare showed lower leaf damage levels (Figure 1). After 12 weeks of infestation, leaves of *O. barthii* were dry and the leaf area was totally damaged. The species *O. glaberrima* presented infestation level 4 (above 80% of the damaged leaf area), while Nipponbare presented level 2 (10% to 20% of damaged leaf area) and red rice presented level 3 (40% to 50% of the damaged leaf area), as shown in Figure 1. Both wild species died before reach the reproductive stage, while both *O. sativa* genotypes were able to complete their life cycle and set seeds. Although the plant height was not negatively affected by mite infestation in any of the tested genotypes/species, tiller number decreased under infested conditions in all genotypes, except for Nipponbare (Figure 2).

**Figure 1.**
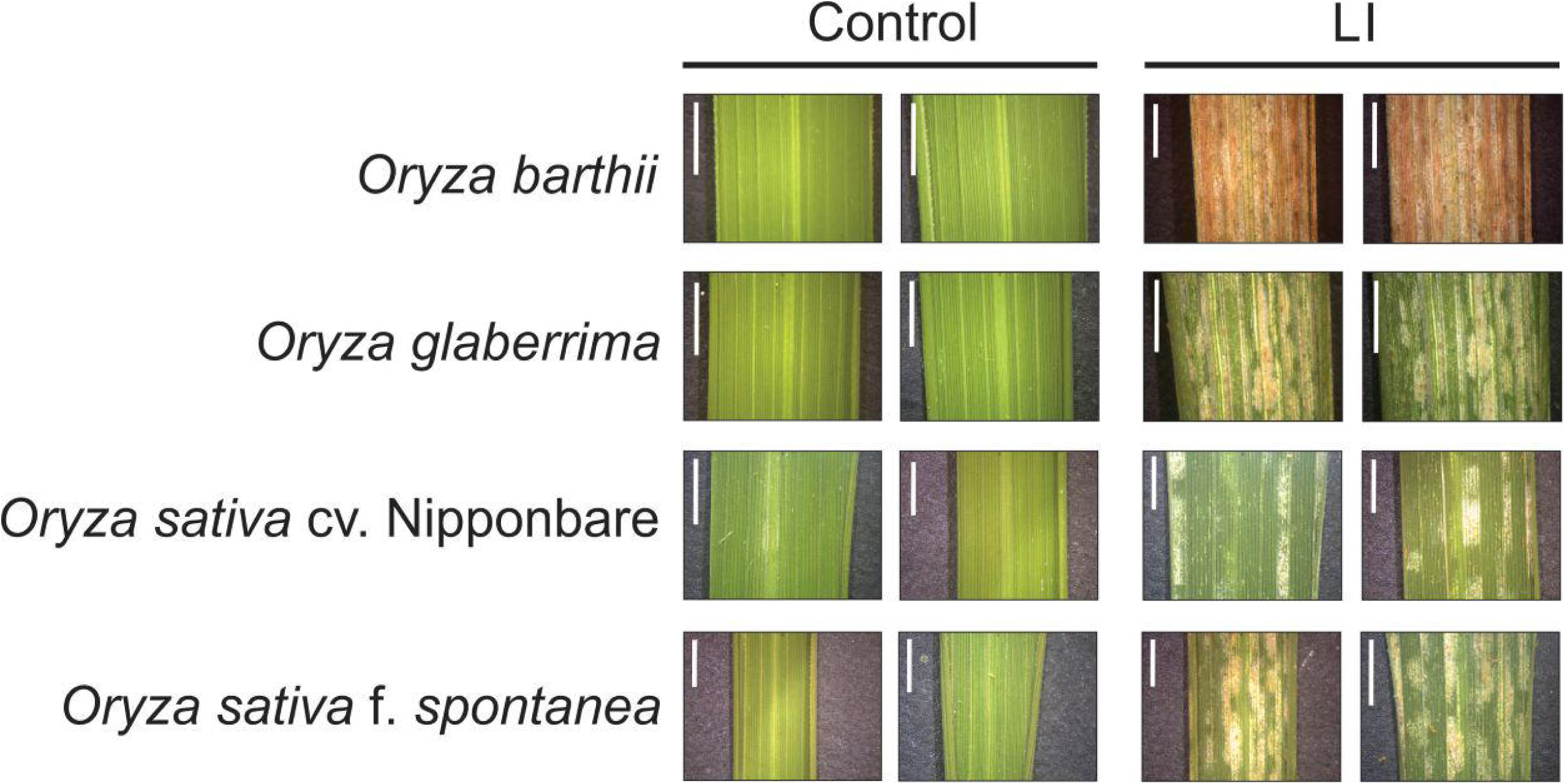
Visual characteristics of leaves from control and late infested (LI) plants of *Oryza barthii*, *Oryza glaberrima*, *Oryza sativa* cv. Nipponbare, and red rice (*Oryza sativa* f. *spontanea*). Bars indicate 0.5 cm.

**Figure 2.**
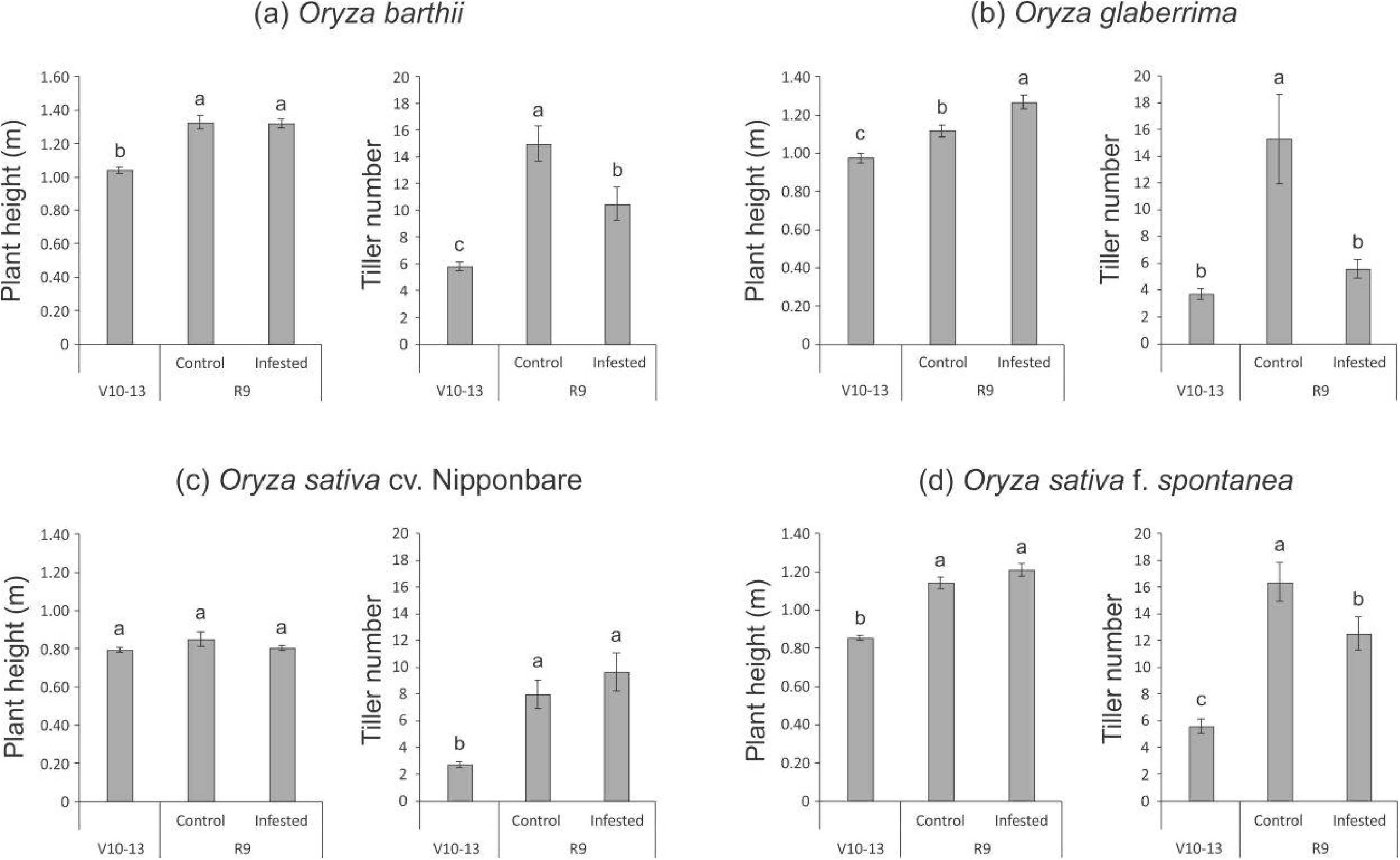
Plant height (m) and tiller number of *Oryza barthii* (a), *Oryza glaberrima* (b), *Oryza sativa* cv. Nipponbare (c), and red rice (*Oryza sativa* f. *spontanea*) (d) at the vegetative stage (no infestation, V10-13) and full maturity stage (control or infested conditions, R9). Represented values are the averages of fifty samples ± SE. Different letters indicate that the means are different by the Tukey HSD test (P ≤ 0.05).

Chlorophyll *a* fluorescence analysis showed that wild rice species (*O. barthii* and *O. glaberrima*) and red rice presented a decrease in at least one of the OJIP curve-times. *O. barthii* seems to be the most affected species, with a decrease in all curve-times after 8 weeks of infestation, while *O. glaberrima* presented a decrease in J-I-P stages, and red rice only in J stage. Nipponbare was the only one with no decrease during the entire OJIP curve under infested condition (Figure 3), suggesting that mite infestation trigger lower damage in Nipponbare photosynthetic apparatus than in the other tested cultivars/species. These results agree with total chlorophyll concentration analysis, where Nipponbare was the only tested cultivars/species with no decrease under infested condition, while *O. barthii* presented the highest decrease comparing infested and control conditions (Figure 4). Also, leaves of the Nipponbare accumulate lower levels of H_2_O_2_ than the other tested cultivars/species, especially when compared to *O. barthii* (Figure 5). Therefore, *S. oryzae* infestation differentially affects the generation of oxidative stress in LI leaves of the tested plants. Altogether, we suggest that *O. barthii* is extremely susceptible to *S. oryzae* infestation, while Nipponbare can be considered tolerant.

**Figure 3.**
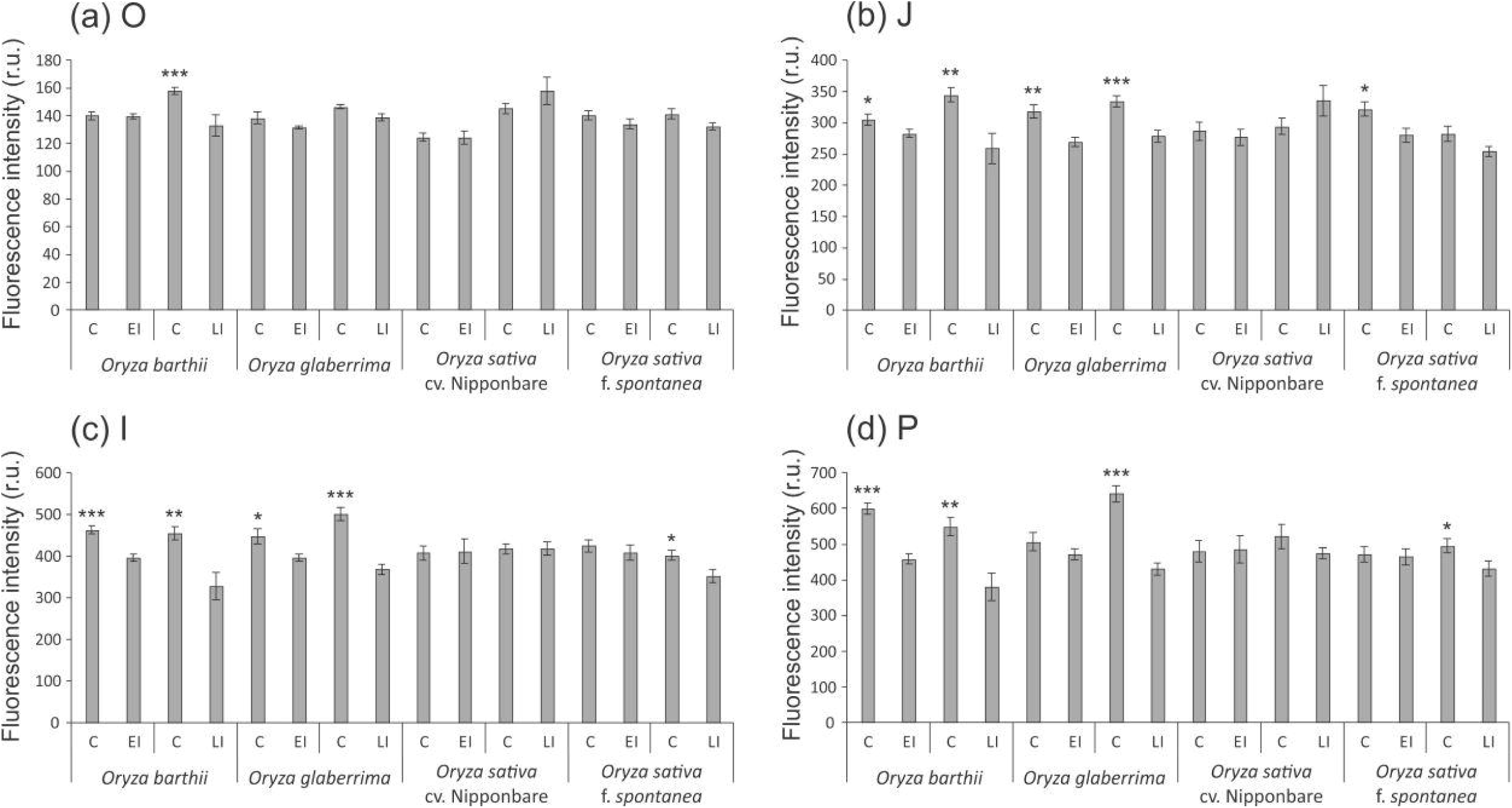
OJIP-test parameters calculated from the chlorophyll *a* fluorescence transient in control (C), early infested (EI), and late infested (LI) leaves of *Oryza barthii*, *Oryza glaberrima*, *Oryza sativa* cv. Nipponbare, and red rice (*Oryza sativa* f. *spontanea*). (a) O; (b) J; (c) I; (d) P. Represented values are the averages of ten samples ± SE. Mean values (from each species/cultivar and each exposure time: C x EI; C x LI) with one, two, or three asterisks are significantly different as determined by a Student’s t test (P ≤ 0.05, 0.01, and 0.001, respectively). r.u. = Raman units.

**Figure 4.**
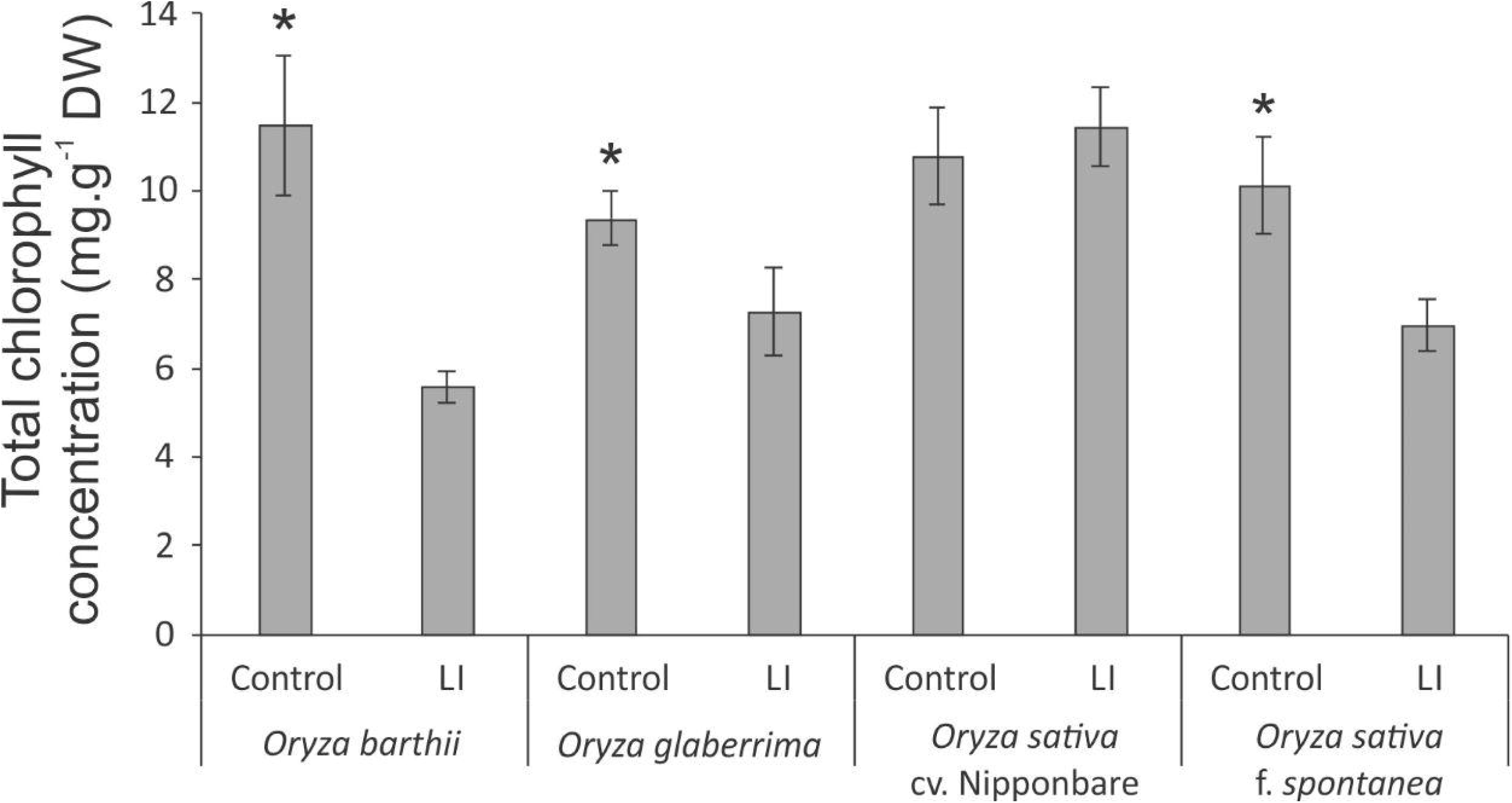
Total chlorophyll concentration in control and late infested (LI) leaves of *Oryza barthii*, *Oryza glaberrima*, *Oryza sativa* cv. Nipponbare, and red rice (*Oryza sativa* f. *spontanea*) plants. Represented values are the averages of three samples ± SE. Mean values (from each species/cultivar: C x LI) with one asterisk are significantly different as determined by a Student’s t test (p-value ≤ 0.05). DW = dry weight.

**Figure 5.**
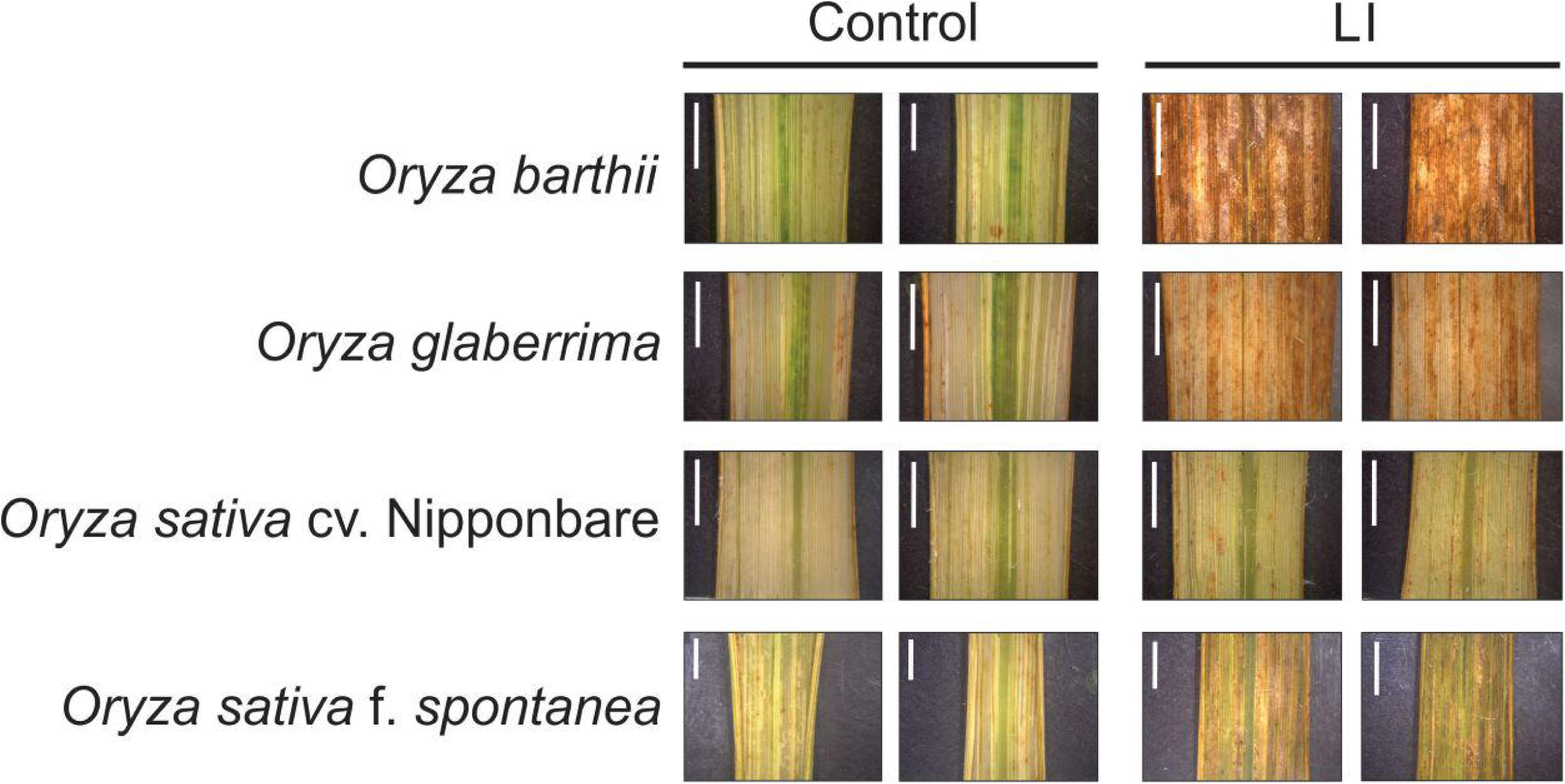
Histochemical staining assay of H_2_O_2_ by diaminobenzidine (DAB) in control and late infested (LI) leaves of *Oryza barthii*, *Oryza glaberrima*, *Oryza sativa* cv. Nipponbare, and red rice (*Oryza sativa* f. *spontanea*). The positive staining (detected in higher levels on infested leaves) in the photomicrographs shows as bright images (brown-color). Bars indicate 0.5 cm.

As wild species were not able to reach the reproductive stage, seeds from Nipponbare were evaluated in order to verify whether *S. oryzae* infestation can impact seed production. As seen in Figure 6, none of the analyzed parameters was negatively affected by mite infestation, and two parameters (percentage of full seeds per panicle and grain length) were even higher under infested condition than under control condition, resulting in the maintenance of seed weight per plant (an estimate of yield) under infested condition (Figure 7). Therefore, Nipponbare presents physiological characteristics of tolerance, since even under infestation, plants can maintain development, photosynthetic and antioxidant activities, and grain yield.

**Figure 6.**
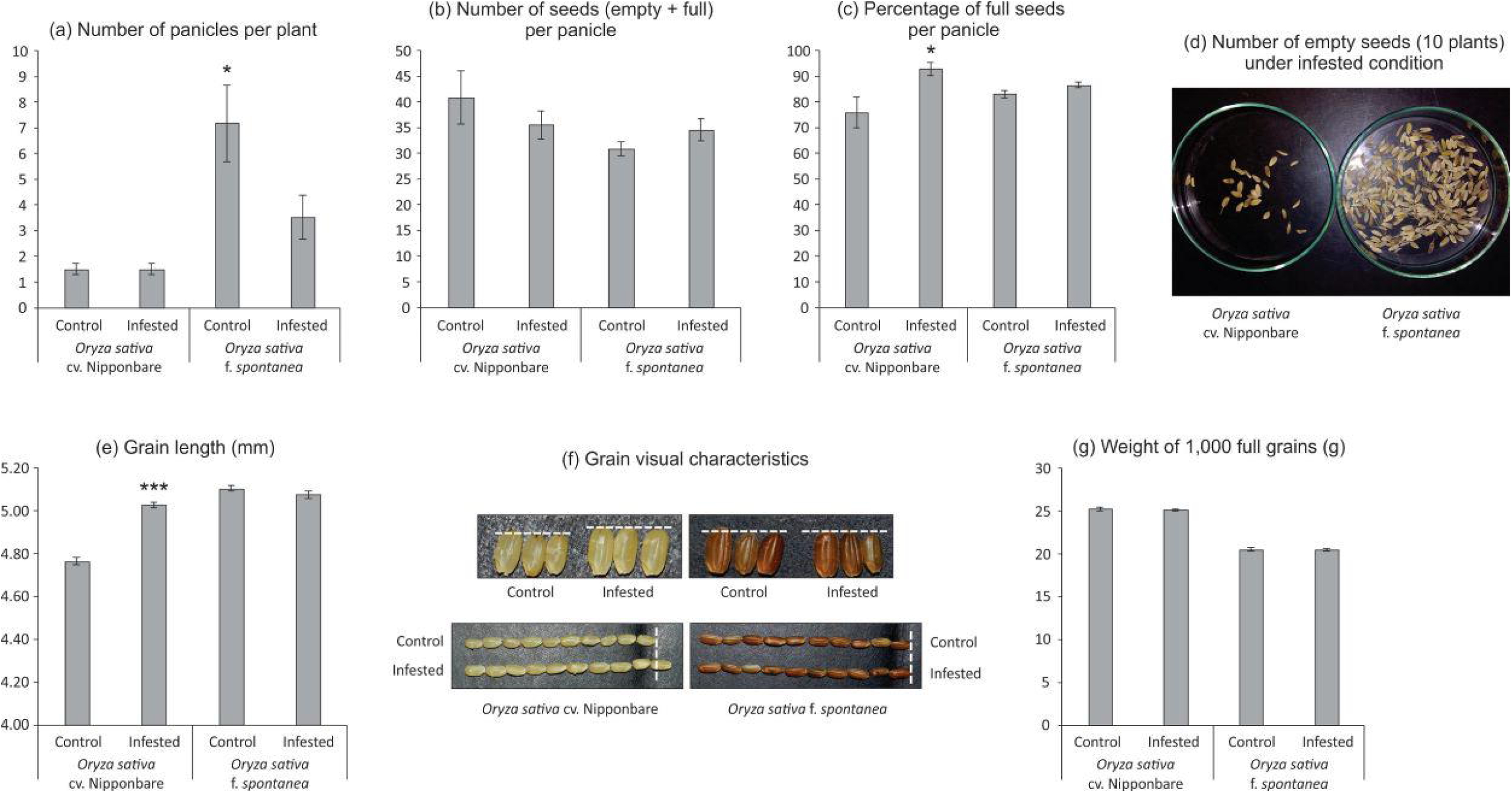
Seeds analysis of *Oryza sativa* cv. Nipponbare and red rice (*Oryza sativa* f. *spontanea*). (a) Number of panicles per plant; (b) Number of seeds (empty + full) per panicle; (c) Percentage of full seeds per panicle; (d) Number of empty seeds (10 plants) under infested condition; (e) Grain length (mm); (f) Grain visual characteristics; and (g) Weight of 1,000 full grains (g). Represented values are the averages of fifty samples ± SE. Mean values (from each cultivar: Control x Infested) with one or three asterisks are significantly different as determined by a Student’s t test (P ≤ 0.05 and 0.001).

**Figure 7.**
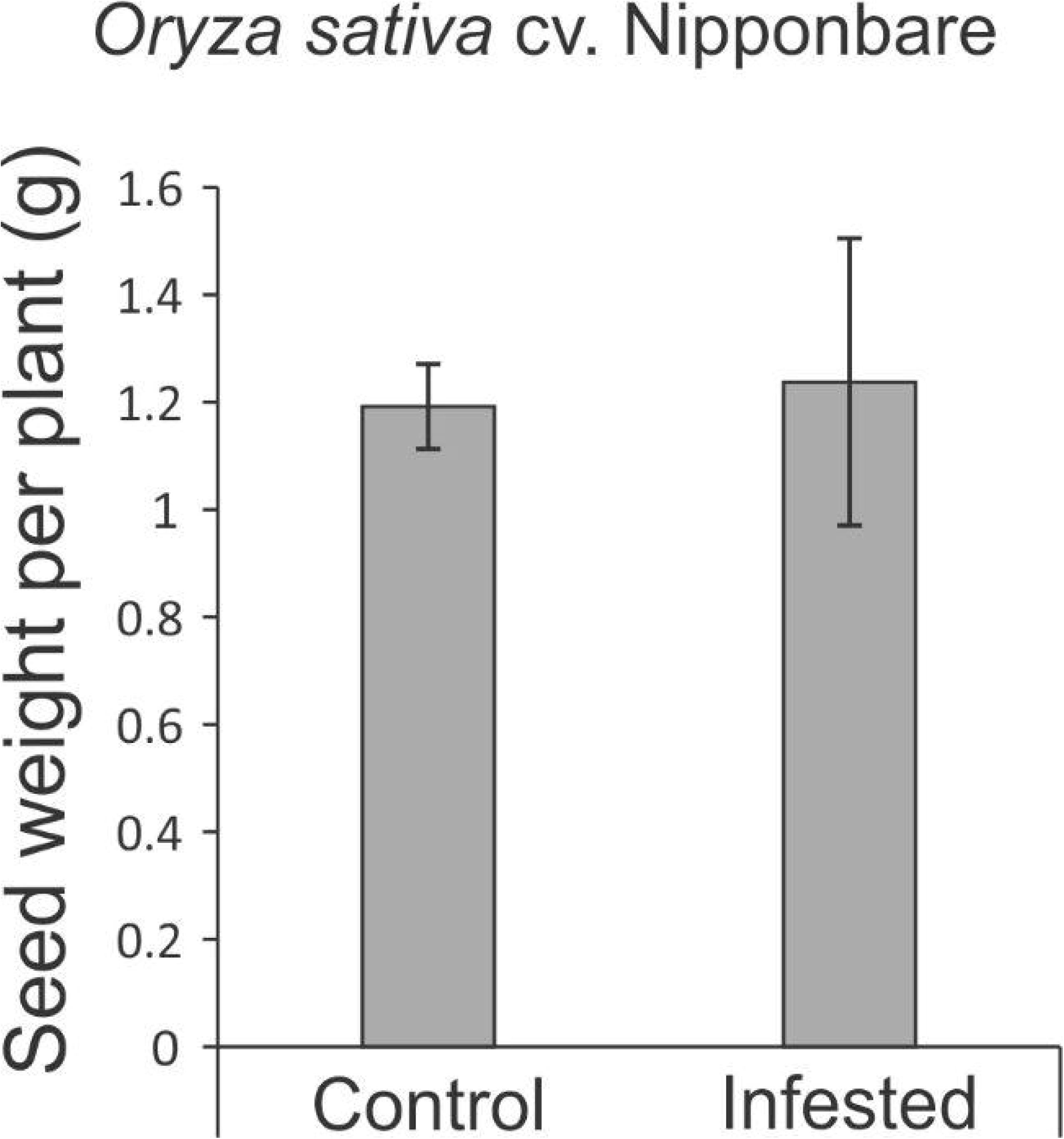
Seed weight per plant (estimative of yield) of *Oryza sativa* cv. Nipponbare under control and infested conditions. Represented values are the averages of ten samples ± SE.

### Overview of proteomic analysis

A total of 1,041 proteins were identified comparing control and infested conditions in both cultivars, with 195 (18.7%) unique to or differentially abundant between treatments. As seen in Figure 8, comparing control and infested leaves of *O. barthii*, we detected 48 proteins, being two more abundant (and one unique) in control condition, and 44 more abundant (and one unique) in infested condition. We identified 147 proteins in control and infested conditions of Nipponbare, being 35 more abundant (and two unique) in control condition, and 104 more abundant (and six unique) in infested condition. Due to the different genetic background between *O. barthii* and *O. sativa*, we prefer not to compare the same condition between species, in order to avoid the emergence of species-specific sequences that are not mite-related.

**Figure 8.**
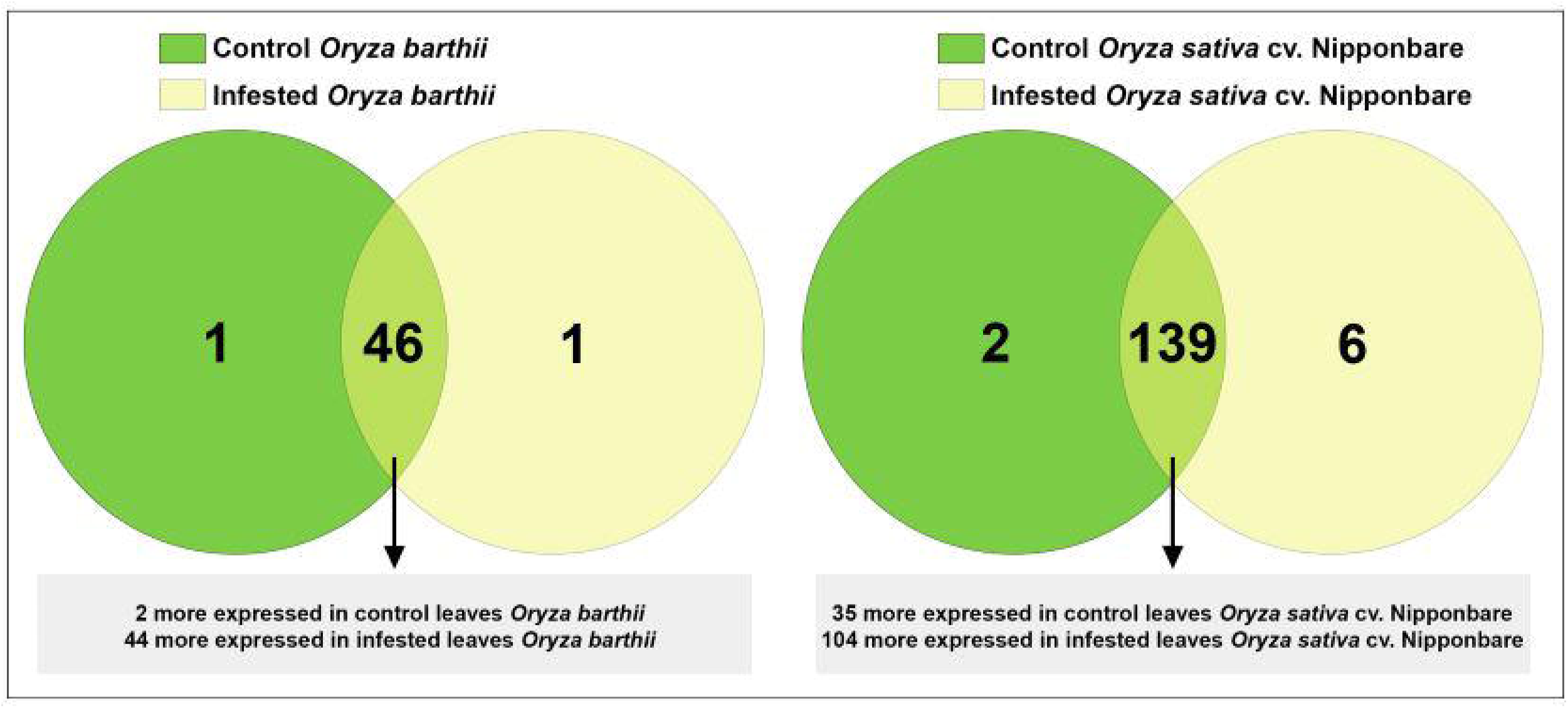
Venn diagram showing the overlap of rice proteins identified in control and early infested (EI) leaves of mite susceptible *Oryza barthii* (a) and mite tolerant *Oryza sativa* cv. Nipponbare (b). Dark green circles: control leaves; yellow circles: EI leaves. Light green means overlap.

The corresponding sequence of each identified protein was compared to NCBI BLASTp to identify specific domains, molecular functions, and protein annotations. Afterward, proteins were categorized in functional categories, according to their putative molecular function. The lists of all unique or differentially abundant proteins identified in this work are presented in Supplementary Tables 1 and 2. In order to facilitate the overall understanding of the differential protein abundance in each functional category presented in Supplementary Tables 1 and 2, we summarize these data in Table 1. We detected four different patterns, based on the number of differentially abundant proteins: 1) most of the functional categories (antioxidant system, carbohydrate metabolism/energy production, general metabolic processes, hormone-related, lipid metabolism, protein modification/degradation, stress response, and others) were more represented in infested condition, regardless the analyzed species; 2) four categories were differentially represented only in one species (amino acid metabolism in *O. barthii* under infested condition; cytoskeleton and transport in Nipponbare under control condition; and protease inhibitor also in Nipponbare, but under infested condition); 3) photosynthesis (over represented in *O. barthii* under infested condition and in Nipponbare under control condition); and 4) translation, with no difference between control and infested condition, regardless the analyzed species.

**Table 1.** Schematic representation of the functional categories differently represented in leaves of mite susceptible *Oryza barthii* and mite tolerant *Oryza sativa* cv. Nipponbare plants under control and infested conditions. Different shades of gray color represent the different patterns of response. Number of arrows (one or two) relates to the representation level, which is based on the number of differentially abundant sequences.

| Functional categories | *Oryza barthii*<br>(susceptible) |  | *Oryza sativa* cv.<br>Nipponbare<br>(tolerant) |  |
| --- | --- | --- | --- | --- |
|  | Control | Infested | Control | Infested |
| Amino acid metabolism | - | ↑ | ↑↑ | ↑↑ |
| Antioxidant system | ↑ | ↑↑ | ↑ | ↑↑ |
| Carbohydrate metabolism/energy production | - | ↑↑ | ↑ | ↑↑ |
| Cytoskeleton | ↑ | ↑ | ↑↑ | - |
| General metabolic processes | - | ↑↑ | ↑ | ↑↑ |
| Hormone-related | - | ↑ | - | ↑↑ |
| Lipid metabolism | - | ↑↑ | - | ↑↑ |
| Photosynthesis | - | ↑↑ | ↑↑ | ↑ |
| Protease inhibitor | - | - | - | ↑↑ |
| Protein modification/degradation | - | ↑↑ | ↑ | ↑↑ |
| Stress response | - | ↑↑ | ↑ | ↑↑ |
| Translation | ↑ | ↑ | ↑↑ | ↑↑ |
| Transport | - | - | ↑↑ | - |
| Others | ↑ | ↑↑ | ↑ | ↑↑ |

(-): functional category not detected as differentially represented.

On the tolerant Nipponbare, *S. oryzae* infestation seems to be less damaging and to generate a more complex defense response. It is interesting to highlight the higher diversity and expression level of antioxidant proteins in Nipponbare than *O. barthii* under infested condition, which agree with the lower oxidative stress seen in Nipponbare leaves (Figure 5), and also the higher diversity of proteins involved with general metabolic processes and carbohydrate metabolism/energy production, suggesting that Nipponbare can maintain the basal/primary metabolism more active than *O. barthii* under infested condition. This also agrees with our observation that photosynthesis is maintained in Nipponbare, whereas it is drastically affected in *O. barthii*. At the same time, Nipponbare seems more able to fight *S. oryzae* infestation than *O. barthii*, presenting higher diversity and/or representation level of protease inhibitors and proteins involved with stress response. Some proteins belonging to these functional categories are discussed later.

## Discussion

According to our first screening of *Oryza* species/genotypes responses to *S. oryzae* infestation, it was shown that none of them present a classical resistance response involving antibiosis and antixenosis (Stenberg and Muola, 2017), since mite population likely increased over time for all species/genotypes, even though with different kinetics (Figure 1). Physiological analysis and agronomical parameters showed that *O. glaberrima* and especially *O. barthii* are extremely susceptible to *S. oryzae* infestation (being unable to reach the reproductive stage), while red rice and especially Nipponbare are less affected by mite infestation (Figure 1 - 7). Recently we detected an *indica* rice cultivar able to maintain seed yield under *S. oryzae* infested condition, which we characterized as mite tolerant (Buffon et al. 2018). Even though wild plant species have been widely recognized as valuable source of resistance genes for developing herbivore-resistant cultivars, Veasey et al. (2008) and Chandrasena et al. (2016) found no signs of mite resistance when different wild rice species were respectively tested against the infestation of *S. oryzae* and *Steneotarsonemus spinki*. These data differ from previously described ones, where *O. barthii* and *O. glaberrima* are recognized as resistant to diseases and pests (Khush, 1997; Linares et al. 2002). Still, our report is one of the first to test mite tolerance in wild rice species.

Recently, Sperotto et al. (2018b) hypothesized that mite-sensitivity presented by wild rice species could be explained, at least partially, by a presumable high Gibberellin (GA):Jasmonic acid (JA) ratio in these plants, since most of the wild rice species are tall plants and probably have high levels of GA synthesis. The roles of GA-JA in growth-defense conflicts during herbivory is not yet fully characterized. However, it is already known that plants need to prioritize GA- or JA-induced responses (Yang et al. 2019). Even though proteins related with GA biosynthesis have not been detected as differentially abundant in *O. barthii* or Nipponbare under infested conditions (Supplementary Table 1 and 2), the height of these plants in full maturity stage is quite different (55% higher in *O. barthii* - Figure 2). It is important to highlight that *O. barthii* and Nipponbare plant heights are not affected by *S. oryzae* infestation (Figure 2), and this could explain the lack of differentially abundant proteins involved with GA biosynthesis when we compare control and infested conditions. Sperotto et al. (2018b) suggest short *Oryza* species (*O. minuta*, *O. meyeriana*, *O. neocaledonica*, and *O. schlechteri*) as primary sources of herbivory tolerance, including mites. According to these authors, plants with low GA level/sensitivity would probably amplify JA-responses and drive plant resources toward defense instead of growth. Based on our data, the GA-based mechanism is not involved in the different tolerance level found in the tested species, although JA might play a role.

One of the most abundant protein found in Nipponbare under infested condition is lipoxygenase (log2 FC 4.7), which is well known as precursor of plant defense mechanisms. Once the herbivore feeds on plant tissues, polyunsaturated fatty acids are released from cell membranes, and accumulate at the wound site (Roach et al. 2015). Also, polyunsaturated fatty acids act as substrate to lipoxygenase, leading to the production of several defense-related compounds, including different structures of JAs such as methyl jasmonate (MeJA) (Mochizuki et al. 2016; Yang et al. 2019). We detected a key enzyme related to JA biosynthesis, allene oxide cyclase (AOC), which converts allene oxide to 12-oxophytodienoic acid (Nguyen et al. 2020), more abundant in both species under infested condition. However, the log2 FC is higher in Nipponbare than in *O. barthii* (Supplementary Table 1 and 2). Therefore, it is possible that Nipponbare presents a more active JA biosynthesis/signaling pathway, which can probably contribute to mite tolerance.

The functional category most affected by *S. oryzae* infestation was antioxidant system, with 10 proteins more abundant in the mite susceptible *O. barthii* (Supplementary Table 1) and 22 in the mite tolerant Nipponbare (Supplementary Table 2) under infested condition. This is an indicative that both species try to defend and maintain its redox homeostasis, since it is governed by the presence of large antioxidant pools that absorb and buffer reductants and oxidants (Foyer and Noctor, 2005). Considering the leaf damage (Figure 1), H_2_O_2_ accumulation (Figure 5), and the high diversity of differentially abundant proteins (Supplementary Tables 1 and 2), such strategy seems to be more efficient in the mite tolerant Nipponbare than in *O. barthii*. Six proteins identified as more abundant under infested condition only in Nipponbare are ferredoxin, ferredoxin-NADP^+^ reductase, thioredoxin H-type, peroxiredoxin, glutathione reductase, and superoxide dismutase (Supplementary Table 2). The stromal ferredoxin-thioredoxin system, a regulatory mechanism linking light to the activity of associated enzymes, is the best-characterized redox signal transduction system in plants, functioning in the regulation of photosynthetic carbon metabolism (Foyer and Noctor, 2005; Schürmann and Buchanan, 2008). Ferredoxin-NADP^+^ reductase catalyzes the reduction of NADP^+^ to NADPH, using the electrons provided by reduced ferredoxin (Cassan et al. 2005). Both NADPH and reduced ferredoxin are required for reductive assimilation and light/dark activation/deactivation of enzymes. Ferredoxin-NADP^+^ reductase is therefore a hub, connecting photosynthetic electron transport to chloroplast redox metabolism (Kozuleva et al. 2016). Even though the precise mechanism remains unknown, a clear correlation between ferredoxin-NADP^+^ reductase content and tolerance to oxidative stress is well established (Giró et al. 2006; Kozuleva et al. 2016).

Thioredoxins H-type are involved in the cellular protection against oxidative stress (Gelhaye et al. 2004), and function as electron donors to several enzymes involved in the protection against oxidative stress such as peroxiredoxin and glutathione reductase (Rouhier et al. 2001). Plants over-expressing thioredoxin H-type genes show oxidative stress tolerance and lower levels of H_2_O_2_ accumulation (Sun et al. 2010; Zhang et al. 2011), similar to our results with the mite tolerant Nipponbare (Figure 5). Peroxiredoxins are abundant low-efficiency peroxidases that act as antioxidant and are involved in modulating redox-dependent signaling cascades (Dietz, 2003). Also, after being reduced by thioredoxin, peroxiredoxins reductively convert H_2_O_2_ to H_2_O at the chloroplast (König et al. 2002), acting together with other peroxidase enzymes that use ascorbate as the electron donor. Ascorbate can be directly regenerated using photosynthetic electrons via reduced ferredoxin or using reduced glutathione (GSH). In the chloroplast, the oxidized glutathione (GSSG) is reduced by the glutathione reductase enzyme using NADPH (Kozuleva et al. 2016). It is important to highlight that a very efficient scavenging mechanisms exist in chloroplasts to prevent oxidative damage, including the rapid conversion of superoxide radical (O_2_^•−^, one of the dominant species produced during light excitation) to H_2_O_2_ by superoxide dismutase enzymes (Mubarakshina and Ivanov, 2010). This is the most likely explanation for the lack of differences in O_2_^•−^ accumulation in the leaves of plants subjected to control and infested conditions (data not shown). Altogether, these data show that the mite tolerant Nipponbare under infested condition can maintain the correct balance between rapid removal of damaging oxidative species and appropriate concentrations necessary to initiate signaling cascades.

We verified that all tested species/genotypes presented a decrease in chlorophyll fluorescence parameters and total chlorophyll concentration, except for the mite tolerant Nipponbare (Figures 3 and 4). Increased photosynthesis is an important mechanism behind tolerance to herbivory (Thomson et al. 2003), and parameters of chlorophyll fluorescence (excess energy absorbed by chlorophyll that is reemitted as light) have been used to obtain quantitative information on the photosynthetic performance of plants (Maxwell and Johnson, 2000), since chlorophyll fluorescence curves allow to evaluate the physiological condition of photosystem II (PSII) components and photosynthetic electron transport chain (Kalaji et al. 2016), where ferredoxin protein acts. Recently, Madriaza et al. (2019) indicated that chlorophyll fluorescence is an accurate predictor of tolerance to herbivore damage and claimed that this rapid methodology could be used to estimate plant tolerance to herbivory in natural populations.

Even though plant height was not affected by *S. oryzae* infestation in *O. barthii* and Nipponbare, the mite susceptible *O. barthii* presented a decrease in tiller number under infested condition, while Nipponbare maintained the same level found in control condition (Figure 2), resulting in no yield losses (Figure 7). Reduction in tillering (and consequently in grain weight) was already seen in rice plants infested by the rice water weevil, *Lissorhoptrus oryzophilus* Kuschel (Zou et al. 2004), and also in IR22 (a rice variety susceptible to the brown planthopper, *Nilaparvata lugens*) (Horgan et al. 2018). It is important to highlight that planthopper damage to IR22 in field cages was severe and plant death occurred in most of IR22 plants, similar to what we found for *O. barthii* and *O. glaberrima* under infested condition. Both tillering and seed filling processes are expensive for the crop energy balance. As seen in Supplementary Tables 1 and 2, we verified a larger diversity of proteins involved with general metabolic processes and carbohydrate metabolism/energy production under infested condition in tolerant Nipponbare than in susceptible *O. barthii*, suggesting that general metabolism and energy production in Nipponbare are less affected by *S. oryzae* infestation than in *O. barthii*.

Two proteins related to general metabolic processes were identified as more abundant under infested condition only in Nipponbare: CBS domain-containing protein (CBSX1) and cysteine synthase (Supplementary Table 2). In Arabidopsis, AtCBSX1 and AtCBX2 positively regulate thioredoxins, which then reduce target proteins, such as peroxiredoxin, which in turn reduce cellular H_2_O_2_ and regulate its level (Yoo et al. 2011). Recently, Kumar et al. (2018) overexpressed a rice Two Cystathionine-β-Synthase Domain-containing Protein (*OsCBSCBSPB4*) in tobacco. Transgenic seedlings were found to exhibit better growth in terms of delayed leaf senescence, profuse root growth and increased biomass in contrast to the wild-type seedlings when subjected to salinity, dehydration, oxidative and extreme temperature treatments, suggesting that OsCBSCBSPB4 is involved in abiotic stress response. As far as we know, this is the first time that a CBS protein is related to biotic stress tolerance in plants. The biosynthesis of cysteine is a limiting step in the production of glutathione, a thiol implicated in various cellular functions, including scavenging of reactive oxygen species and resistance to biotic stresses (Youssefian et al. 2001). At the same time, several studies have already pointed out the function of cysteine-thiols and sulphur-based primary and secondary metabolites in basal defense responses to abiotic and biotic stresses (for a comprehensive review see Tahir and Dijkwel (2016)). Even though herbivory resistance can employ metabolites containing cysteine such as glucosinolates (Guo et al. 2013), to the best of our knowledge this is the first work that links the expression of cysteine synthase protein with mite tolerance.

Two proteins related with stress response were also identified as more abundant under infested condition only in Nipponbare: osmotin-like protein and ricin B-like lectin. The osmotin protein belongs to the PR-5 family of Pathogenesis-related (PR) proteins, which are produced in response to diseases caused by various biotic and abiotic stresses. Osmotin can inhibit the activity of defensive cell wall barriers and increases its own cytotoxic efficiency, triggering changes in the cell wall and enabling osmotin penetration into the plasma membrane (Hakim et al. 2018). Overexpression of genes encoding osmotins confers tolerance against biotic and abiotic stresses in different plant species (Barthakur et al. 2001; Xue et al. 2016; Chowdhury et al. 2017), besides stimulating the accumulation of proline, which functions as a compatible osmolyte useful in responding to abiotic stress, especially drought and salinity (Hakim et al. 2018). Proline also function suppressing free radicals and preventing chlorophyll degradation in vegetative cells, in order to maintain the photosynthetic processes (Barthakur et al. 2001; Chowdhury et al. 2017). In the mite tolerant Nipponbare we detected higher levels of proline accumulation under infested condition when compared to control (Figure 9), along with maintenance of chlorophyll concentration (Figure 4) after mite infestation. This is the first time that expression of an osmotin protein is related to mite tolerance. Since the generation of transgenic plants overexpressing osmotin genes is being recommended as a biotechnological tool to generate stress tolerance, we are currently overexpressing *OsOSM1* gene (LOC_Os12g38170) in rice plants, along with CRISPR-Cas9 gene knockout.

**Figure 9.**
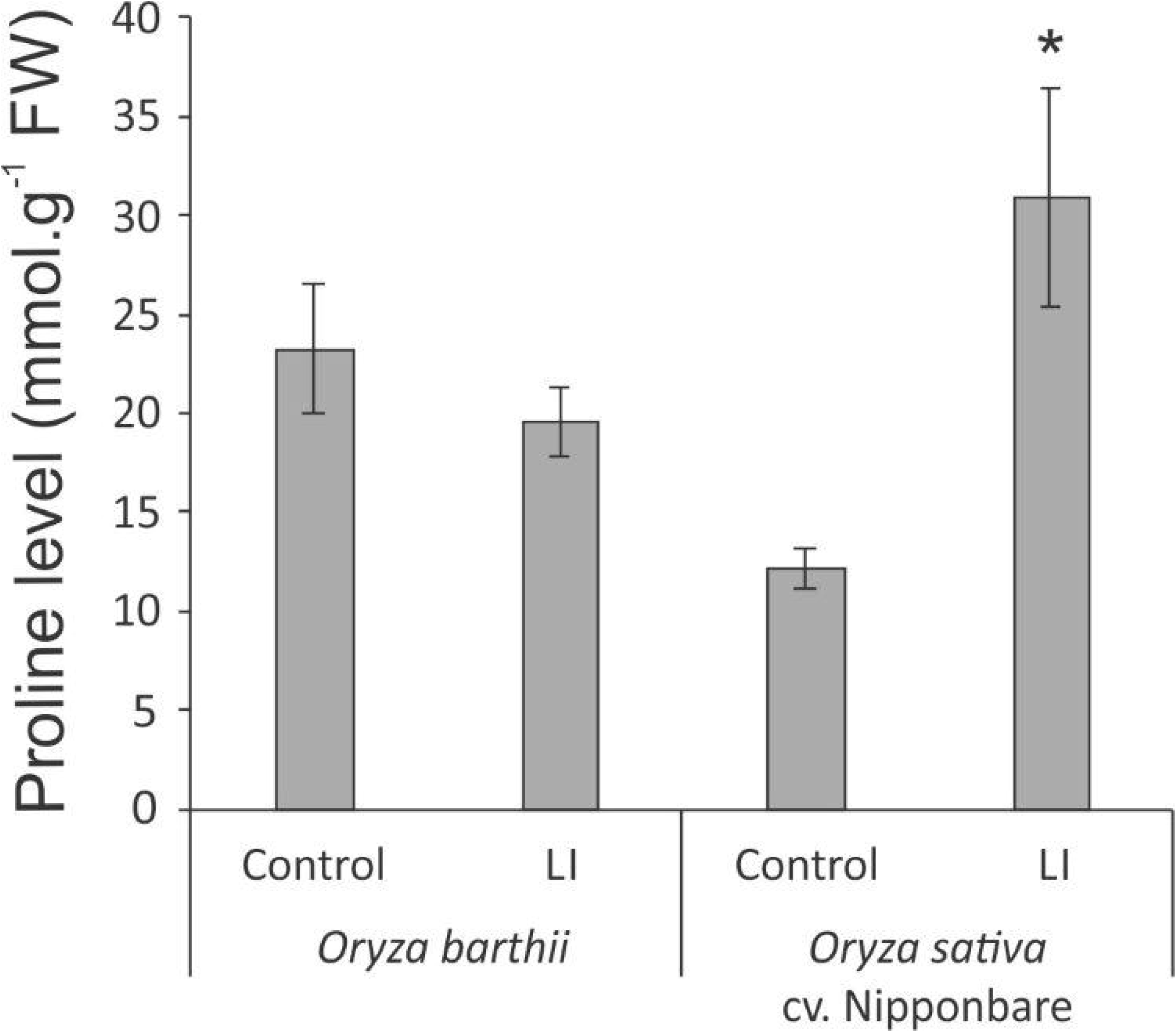
Proline levels in control and late infested (LI) leaves of mite susceptible *Oryza barthii* and mite tolerant *Oryza sativa* cv. Nipponbare. Mean values with one asterisk are significantly different as determined by a Student’s t test (P ≤ 0.05). FW = fresh weight.

Ricin B-like lectins, one of the most widespread families of carbohydrate binding proteins (Lannoo and Van Damme, 2014), have been suggested to play a role in plant defense against pathogens (Vandenbussche et al. 2004) and insects (Wei et al. 2004; Shahidi-Noghabi et al. 2009). Since mutation of the carbohydrate binding site can abolish or reduce the toxic effect, the entomotoxic properties of the proteins can be linked to their carbohydrate binding activity (Shahidi-Noghabi et al. 2008). Our work is the first one that correlates a ricin B-like lectin expression with mite response/tolerance in plants. Curiously, it was shown that in β-trefoil structure enables interactions between fungal ricin B-like lectins and protease inhibitors, and that these interactions modulate their biological activity (Žurga et al. 2015). It would be interesting to test whether the two protease inhibitors (putative Bowman Birk trypsin inhibitor) that we found more abundant only in mite tolerant Nipponbare under infested condition (Supplementary Table 2) are able to interact with the ricin B-like lectins.

Among plant defenses, protease inhibitors (PIs) have direct effects on herbivores, interfering with their physiology (Diaz and Santamaria, 2012; Martinez et al. 2016; Arnaiz et al. 2018). Once ingested by the arthropod, PIs inhibit proteolytic activities, preventing or hindering protein degradation (Terra and Ferreira, 2005). The first successfully overexpressed Bowman-Birk family IP gene in tobacco was the cowpea trypsin inhibitor (*CpTI* - Hilder et al. 1987). After, the *CpTI* gene was inserted into the genome of several plants, increasing the resistance to different herbivore species (Santamaria et al. 2012). Since then, different IPs have been obtained from cultivated plants such as rice, barley, soybean, sweet potato and corn, and have been overexpressed in different plant species, conferring resistance to various pest insects (Clemente et al. 2019). Rice has seven Bowman Birk trypsin inhibitor genes, and overexpression of *RBBI2-3* in transgenic rice plants resulted in resistance to the fungal pathogen *Pyricularia oryzae*, indicating that proteinase inhibitors confer *in vivo* resistance against the fungal pathogen and that they might play a role in the defense system of the rice plant (Qu et al. 2003). Recently, Arnaiz et al. (2018) verified that Kunitz trypsin inhibitors (KTI) participate in the plant defense responses against the spider mite *Tetranychus urticae*. Thus, our work is the first to suggest that Bowman Birk trypsin inhibitors may be involved in the tolerance of rice plants against *S. oryzae* mite infestation.

## Concluding remarks

Tolerance is a sustainable pest management strategy as it only involves plant response, and therefore does not cause evolution of resistance in target pest populations (Peterson et al. 2017; Sperotto et al. 2018a). According to Gerszberg and Hnatuszko-Konka (2017), elucidation of the tolerance mechanism at the biochemical, physiological and morphological level remains one of the greatest challenges of the contemporary plant physiology. Our physiological results showed that wild rice (*O. barthii* and *O. glaberrima*) and red rice (*O. sativa* f. spontanea) do not present tolerance to *S. oryzae* mite infestation. Still, we characterized *O. sativa* cv. Nipponbare as tolerant to mite infestation. Proteomic data showed that *O. barthii* presents a less diverse and efficient antioxidant apparatus under infested condition, not being able to modulate proteins involved with general metabolic processes and energy production, causing damage to its photosynthetic apparatus and development. On the other hand, Nipponbare presents more efficient antioxidant mechanisms that maintain cellular homeostasis and its photosynthetic capacity. Tolerant plants are able to modulate proteins involved with general metabolic processes and energy production, maintaining their development. In addition, these plants can modulate the expression of defense proteins, such as osmotin, ricin B-like lectin, and protease inhibitor, which may be used in future breeding programs for increasing rice tolerance to mite infestation. The model in Figure 10 summarizes the rice tolerance mechanisms putatively employed by Nipponbare.

**Figure 10.**
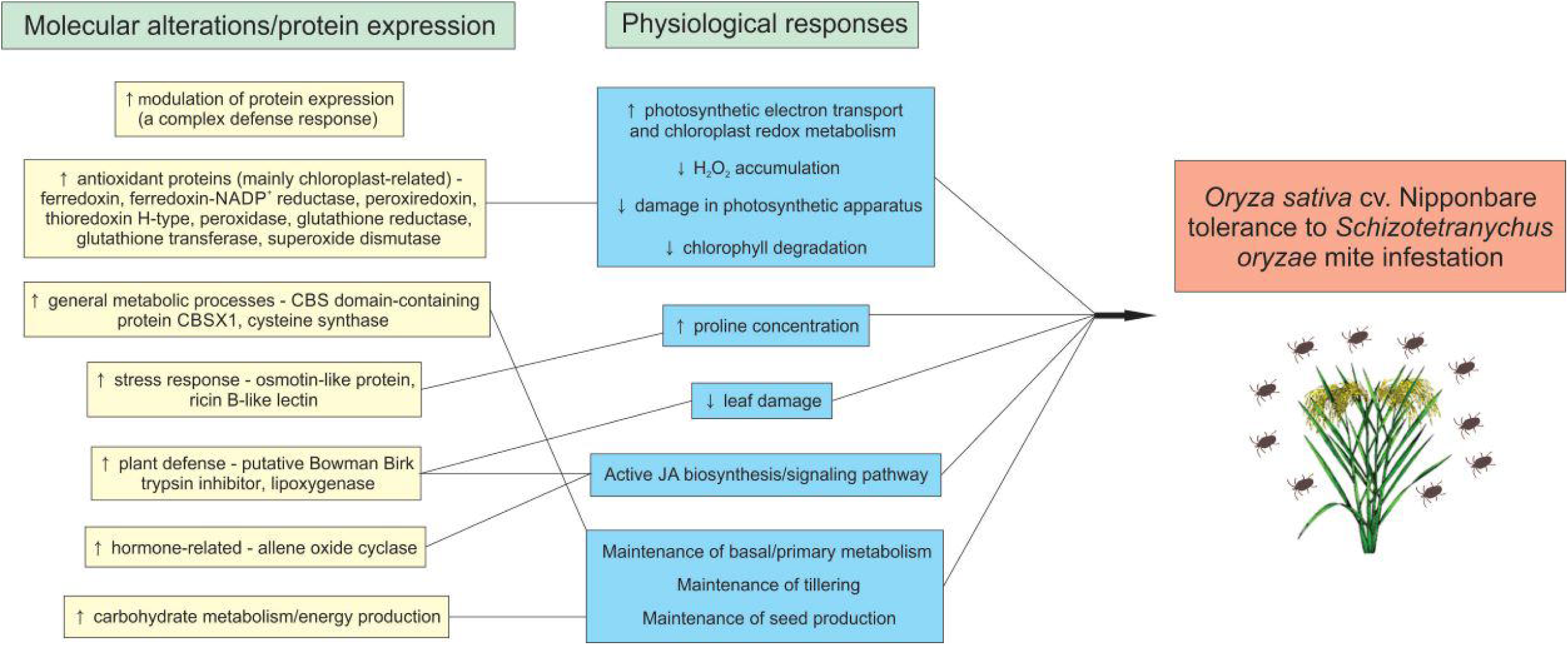
Rice mechanisms employed by *Oryza sativa* cv. Nipponbare to tolerate *Schizotetranychus oryzae* mite infestation.

## Acknowledgments

This research was supported by University of Taquari Valley - Univates, Fundação de Amparo à Pesquisa do Estado do RS (FAPERGS) and Conselho Nacional de Desenvolvimento Científico e Tecnológico (CNPq). The authors thank Instituto Rio-Grandense do Arroz (IRGA) for technical support.

## Author Contributions Statement

JS, FKR, and RAS conceived and designed research. GB, EARB, TIL, JMA, and AB conducted experiments. VS and MCBL contributed with analytical tools. GB, EARB, JMA, AB, VS, and RAS analyzed the data. GB and RAS wrote the manuscript. All authors read and approved the manuscript.

## Supporting Information Available

**Supplementary Table 1.** Differentially abundant proteins in mite susceptible *Oryza barthii* (control x early infested condition).

**Supplementary Table 2.** Differentially abundant proteins in mite tolerant *Oryza sativa* cv. Nipponbare (control x early infested condition).

**Supplementary Figure 1.** Classification of infestation levels according to visual characteristics of leaves.

