## Supplementary figures and images for "Nipponbare and wild rice species as unexpected tolerance and susceptibility sources against *Schizotetranychus oryzae* (Acari: Tetranychidae) mite infestation"

### Supplementary Figure 1

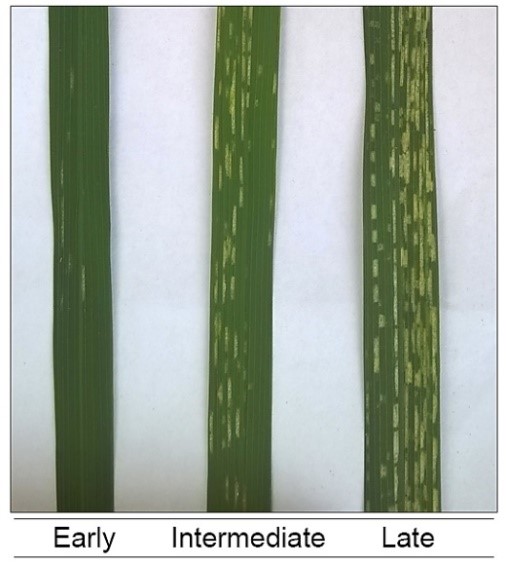
