## Supplementary Table 1 for "Nipponbare and wild rice species as unexpected tolerance and susceptibility sources against *Schizotetranychus oryzae* (Acari: Tetranychidae) mite infestation"

**Supplementary Table 1.** Differentially abundant proteins in mite susceptible *Oryza barthii* (control x early infested condition).

| ***Oryza barthii* Control x Early infested - Proteins unique or more expressed in control leaves** | | | | | | |
| --- | --- | --- | --- | --- | --- | --- |
| **Functional categories** | **Description** | **Uniprot** | **Locus** | **Reported peptides** | **t-Test** | **Log2 Fold Change** |
| **Antioxidant system** | ferredoxin--nitrite reductase, chloroplastic | Q0JMV6_ORYSJ | LOC_Os01g25484 | 11 | 0.046 | -0.825 |
| **Translation** | 60S ribosomal protein L6 | Q6YY64_ORYSJ | LOC_Os02g37862 | 2 | 0.018 | -0.747 |
| **Cytoskeleton** | actin-7 | Q94DL4_ORYSJ | LOC_Os01g73310 | 11 | Unique Control | Unique Control |
| ***Oryza barthii* Control x Early infested - Proteins unique or more expressed in early infested leaves** | | | | | | |
| **Functional categories** | **Description** | **Uniprot** | **Locus** | **Reported peptides** | **t-Test** | **Log2 Fold Change** |
| **Antioxidant system** | peroxidase 2 | Q7F1U0_ORYSJ | LOC_Os07g48020 | 7 | 0.003 | 1.561 |
|  | peroxidase 2-like | Q7F1U1_ORYSJ | LOC_Os07g48010 | 2 | 0.004 | 1.477 |
|  | glutathione transferase GST 23 | Q93WY5_ORYSJ | LOC_Os09g29200 | 3 | 0.018 | 1.307 |
|  | peroxidase 1 | Q654S0_ORYSJ | LOC_Os01g22230 | 4 | 0.019 | 0.940 |
|  | cationic peroxidase SPC4-like | Q5JMS4_ORYSJ | LOC_Os01g73170 | 5 | 0.017 | 0.838 |
|  | glutathione S-transferase 1 | GSTF2_ORYSJ | LOC_Os01g55830 | 4 | 0.012 | 0.821 |
|  | anionic peroxidase | Q7XSV2_ORYSJ | LOC_Os04g59150 | 3 | 0.038 | 0.779 |
|  | lactoylglutathione lyase | LGUL_ORYSJ | LOC_Os08g09250 | 10 | 0.022 | 0.764 |
|  | catalase | CATA2_ORYSJ | LOC_Os06g51150 | 7 | 0.047 | 0.574 |
|  | monodehydroascorbate reductase | MDAR3_ORYSJ | LOC_Os09g39380 | 5 | 0.046 | 0.535 |
| **Stress response** | momilactone A synthase-like | Q0D3V0_ORYSJ | LOC_Os07g46930 | 6 | 0.009 | 1.638 |
|  | momilactone A synthase-like | A0A0P0X9T1_ORYSJ | LOC_Os07g46830 | 2 | 0.031 | 1.250 |
|  | cysteine-rich repeat secretory protein 55-like | Q75L18_ORYSJ | LOC_Os05g02200 | 2 | 0.002 | 1.179 |
|  | PLAT domain-containing protein 3 | Q7XRE7_ORYSJ | LOC_Os04g38390 | 4 | 0.013 | 1.126 |
|  | heat shock cognate 70 kDa protein | Q10NA1_ORYSJ | LOC_Os03g16920 | 10 | 0.003 | 1.022 |
|  | stress-response A/B barrel domain-containing protein UP3 | A0A0P0X8Q0_ORYSJ | LOC_Os07g41820 | 2 | 0.049 | 0.694 |
|  | chalcone--flavonone isomerase | CFI_ORYSJ | LOC_Os03g60509 | 5 | 0.001 | 0.694 |
|  | Grx_C2.2 - glutaredoxin subgroup I | GRXC6_ORYSJ | LOC_Os04g42930 | 2 | 0.038 | 0.527 |
| **Protein modification/degradation** | probable carboxylesterase 15 | Q8GSJ3_ORYSJ | LOC_Os07g06830 | 4 | 0.011 | 1.082 |
|  | serine carboxypeptidase 2 | Q5W6C6_ORYSJ | LOC_Os05g18604 | 2 | 0.003 | 0.903 |
|  | serine carboxypeptidase II-2 | Q5SMV5_ORYSJ | LOC_Os06g08720 | 2 | 0.030 | 0.861 |
|  | oryzain alpha chain | ORYA_ORYSJ | LOC_Os04g55650 | 4 | 0.000 | 0.806 |
|  | subtilisin-like protease SBT1.9 | Q8S1N3_ORYSJ | LOC_Os01g64860 | 5 | 0.017 | 0.748 |
|  | probable carboxylesterase 15 | Q8GSE8_ORYSJ | LOC_Os07g06840 | 4 | 0.018 | 0.647 |
|  | serine carboxypeptidase-like 50 | Q75HY2_ORYSJ | LOC_Os05g50600 | 7 | 0.037 | 0.640 |
| **General metabolic processes** | putative Tryptophan synthase beta chain | Q67VM2_ORYSJ | LOC_Os06g42560 | 3 | 0.028 | 1.049 |
|  | primary amine oxidase | A0A0P0W896_ORYSJ | LOC_Os04g20164 | 2 | 0.049 | 0.697 |
|  | nucleoside diphosphate kinase 2, chloroplastic | A0A0P0YB67_ORYSJ | LOC_Os12g36194 | 2 | 0.010 | 0.620 |
|  | glyceraldehyde-3-phosphate dehydrogenase 1, cytosolic | G3PC1_ORYSJ | LOC_Os08g03290 | 13 | 0.019 | 0.537 |
| **Lipid metabolism** | lipid-transfer protein | NLTP1_ORYSJ | LOC_Os12g02320 | 2 | 0.015 | 2.117 |
|  | lipid transfer protein | NLT2A_ORYSJ | LOC_Os11g02369 | 4 | 0.043 | 1.785 |
|  | 3-ketoacyl-CoA thiolase 2, peroxisomal-like | Q84P96_ORYSJ | LOC_Os02g57260 | 6 | 0.002 | 0.576 |
| **Carbohydrate metabolism and energy production** | NADP-dependent malic enzyme isoform X1 | A0A0P0V7M3_ORYSJ | LOC_Os01g52500 | 6 | 0.016 | 0.798 |
|  | thiamine thiazole synthase 2, chloroplastic | A0A0N7KNK4_ORYSJ | LOC_Os07g34570 | 4 | 0.014 | 0.691 |
|  | transaldolase 2 | Q5JK10_ORYSJ | LOC_Os01g70170 | 13 | 0.029 | 0.538 |
| **Photosynthesis** | protochlorophyllide reductase A, chloroplastic | PORA_ORYSJ | LOC_Os04g58200 | 4 | 0.032 | 0.923 |
|  | ruBisCO large subunit-binding protein subunit beta, chloroplastic | Q6ZFJ9_ORYSJ | LOC_Os02g01280 | 5 | 0.004 | 0.717 |
| **Hormone-related** | allene oxide cyclase, chloroplastic | AOC_ORYSJ | LOC_Os03g32314 | 3 | 0.032 | 0.523 |
| **Amino acid metabolism** | peptide methionine sulfoxide reductase A4, chloroplastic | MSRA4_ORYSJ | LOC_Os10g41400 | 2 | 0.042 | 0.511 |
| **Translation** | putative RNase S-like protein precursor | Q69JF3_ORYSJ | LOC_Os09g36700 | 3 | 0.045 | 1.169 |
| **Cytoskeleton** | actin-2 | ACT2_ORYSJ | LOC_Os10g36650 | 13 | Unique Infected | Unique Infected |
|  | chitin elicitor-binding protein-like | CEBIP_ORYSJ | LOC_Os03g04110 | 3 | 0.001 | 0.958 |
| **Others** | amidase 1 | A3AQC6_ORYSJ | LOC_Os04g02754 | 5 | 0.008 | 0.910 |
|  | PLAT domain-containing protein 3 | Q6ZGP5_ORYSJ | LOC_Os02g51710 | 5 | 0.043 | 0.765 |
|  | dirigent protein 21 | Q2R0H3_ORYSJ | LOC_Os11g42550 | 5 | 0.028 | 0.684 |
