## Supplementary Table 2 for "Nipponbare and wild rice species as unexpected tolerance and susceptibility sources against *Schizotetranychus oryzae* (Acari: Tetranychidae) mite infestation"

**Supplementary Table 2.** Differentially abundant proteins in mite tolerant *Oryza sativa* cv. Nipponbare (control x early infested condition).

| ***Oryza sativa* cv. Nipponbare Control x Early infested - Proteins unique or more expressed in control leaves** | | | | | | |
| --- | --- | --- | --- | --- | --- | --- |
| **Functional categories** | **Description** | **Uniprot** | **Locus** | **Reported peptides** | **t-Test** | **Log2 Fold Change** |
| **Photosynthesis** | photosystem II 47 kDa protein (chloroplast) | PSBB_ORYSJ | LOC_Os06g39708 | 8 | 0.004 | -1.672 |
|  | carboxyvinyl-carboxyphosphonate phosphorylmutase, chloroplastic | Q0IPL3_ORYSJ | LOC_Os12g08760 | 5 | 0.002 | -1.292 |
|  | photosystem II protein D2 (chloroplast) | PSBD_ORYSJ | LOC_Osp1g00170 | 5 | 0.018 | -1.253 |
|  | photosystem II CP43 chlorophyll apoprotein (chloroplast) | PSBC_ORYSJ | LOC_Osp1g00180 | 7 | 0.024 | -1.198 |
|  | protochlorophyllide reductase B, chloroplastic | PORB_ORYSJ | LOC_Os10g35370 | 2 | 0.000 | -1.077 |
|  | phosphoenolpyruvate carboxylase | Q5QNA5_ORYSJ | LOC_Os01g11054 | 2 | 0.001 | -0.998 |
|  | photosystem II protein D1 (chloroplast) | PSBA_ORYSJ | LOC_Osp1g00110 | 3 | 0.014 | -0.993 |
|  | protein TIC110, chloroplastic | Q0IWS0_ORYSJ | LOC_Os10g35030 | 4 | 0.046 | -0.976 |
|  | transketolase, chloroplastic | Q7XWP9_ORYSJ | LOC_Os04g19740 | 3 | 0.001 | -0.764 |
|  | chlorophyll a-b binding protein 7, chloroplastic | Q6ZL95_ORYSJ | LOC_Os07g38960 | 5 | 0.011 | -0.761 |
|  | chlorophyll a-b binding protein CP29.1, chloroplastic | Q6Z411_ORYSJ | LOC_Os07g37240 | 3 | 0.007 | -0.739 |
|  | chlorophyll a-b binding protein of LHCII type III, chloroplastic | Q6ZF30_ORYSJ | LOC_Os07g37550 | 8 | 0.017 | -0.537 |
| **Translation** | 50S ribosomal protein L11, chloroplastic | A0A0P0VSE5_ORYSJ | LOC_Os03g03020 | 2 | 0.012 | -1.109 |
|  | ribosomal protein L2 (chloroplast) | RK2_ORYSJ | LOC_Osp1g01100 | 3 | 0.000 | -0.910 |
|  | ribosomal protein L16 (chloroplast) | RK16_ORYSJ | LOC_Os05g22724 | 3 | 0.000 | -0.879 |
|  | elongation factor G-2, chloroplastic | A0A0N7KJF4_ORYSJ | LOC_Os04g45490 | 4 | 0.000 | -0.792 |
|  | 50S ribosomal protein L4, chloroplastic | Q10NM5_ORYSJ | LOC_Os03g15870 | 2 | 0.001 | -0.722 |
|  | 30S ribosomal protein S1, chloroplastic | Q0DSD6_ORYSJ | LOC_Os03g20100 | 9 | 0.018 | -0.516 |
| **Cytoskeleton** | tubulin alpha-1 chain | TBA2_ORYSJ | LOC_Os11g14220 | 15 | 0.003 | -1.170 |
|  | tubulin beta-5 chain | TBB1_ORYSJ | LOC_Os01g18050 | 16 | 0.005 | -0.100 |
|  | tubulin alpha-1 chain | TBA1_ORYSJ | LOC_Os07g38730 | 13 | 0.017 | -0.656 |
|  | tubulin beta chain | TBB7_ORYSJ | LOC_Os03g56810 | 17 | 0.014 | -0.650 |
|  | tubulin beta chain | TBB3_ORYSJ | LOC_Os06g46000 | 17 | 0.020 | -0.581 |
| **Carbohydrate metabolism and energy production** | sucrose synthase 3 | SUS3_ORYSJ | LOC_Os07g42490 | 11 | Unique Control | Unique Control |
|  | ATP synthase CF0 subunit I (chloroplast) | ATPF_ORYSJ | LOC_Os10g38272 | 2 | 0.021 | -1.137 |
|  | sucrose synthase 1 | SUS1_ORYSJ | LOC_Os03g28330 | 26 | 0.001 | -0.687 |
| **Antioxidant system** | ferredoxin--nitrite reductase, chloroplastic | NIR_ORYSJ | LOC_Os01g25484 | 11 | 0.000 | -1.872 |
|  | peroxidase 70 | Q5U1Q2_ORYSJ | LOC_Os03g22010 | 4 | 0.019 | -1.480 |
|  | peroxidase 1-like | Q0DCP0_ORYSJ | LOC_Os06g20150 | 7 | 0.028 | -0.885 |
| **Amino acid metabolism** | glycine dehydrogenase (decarboxylating), mitochondrial | A0A0P0V7A3_ORYSJ | LOC_Os01g51410 | 11 | 0.009 | -0.717 |
|  | glycine dehydrogenase (decarboxylating), mitochondrial | A0A0P0WYZ5_ORYSJ | LOC_Os06g40940 | 18 | 0.005 | -0.659 |
| **General metabolic processes** | probable pyridoxal 5'-phosphate synthase subunit PDX1.1 | Q53NW9_ORYSJ | LOC_Os11g48080 | 5 | 0.004 | -0.669 |
| **Protein modification/degradation** | serine/threonine-protein phosphatase 2A 65 kDa regulatory subunit A beta isoform X1 | A0A0P0XJR1_ORYSJ | LOC_Os09g07510 | 3 | 0.002 | -0.600 |
| **Transport** | Aquaporin PIP1-2 | PIP12_ORYSJ | LOC_Os04g47220 | 4 | 0.006 | -1.666 |
|  | putative cellular retinaldehyde-binding/triple function | Q5TKJ2_ORYSJ | LOC_Os05g35460 | 6 | 0.004 | -0.834 |
| **Stress response** | heat shock protein 90-5, chloroplastic | Q6ZCV7_ORYSJ | LOC_Os08g38086 | 4 | 0.035 | -0.545 |
| **Others** | putative beta-D-glucan exohydrolase | Q6ZG89_ORYSJ | LOC_Os02g03870 | 2 | Unique Control | Unique Control |
| ***Oryza sativa* cv. Nipponbare Control x Early infested - Proteins unique or more expressed in early infested leaves** | | | | | | |
| **Functional categories** | **Description** | **Uniprot** | **Locus** | **Reported peptides** | **t-Test** | **Log2 Fold Change** |
| **Antioxidant system** | peroxidase 2 | PER2_ORYSJ | LOC_Os07g48030 | 3 | Unique Infested | Unique Infested |
|  | GDP-mannose 3,5-epimerase 2 | GME2_ORYSJ | LOC_Os11g37890 | 4 | Unique Infested | Unique Infested |
|  | peroxidase P7-like | Q5Z7J2_ORYSJ | LOC_Os06g35520 | 2 | 0.026 | 3.482 |
|  | probable glutathione S-transferase GSTU6 | GSTU6_ORYSJ | LOC_Os10g38740 | 5 | 0.015 | 2.430 |
|  | peroxidase BP1 precursor [*Oryza nivara*] | Q94DM2_ORYSJ | LOC_Os01g73200 | 5 | 0.001 | 1.926 |
|  | glutathione transferase GST 23 | Q93WY5_ORYSJ | LOC_Os09g29200 | 4 | 0.001 | 1.884 |
|  | peroxidase 2 | Q7F1U0_ORYSJ | LOC_Os07g48020 | 7 | 0.005 | 1.833 |
|  | probable L-gulonolactone oxidase 4 | Q6YXT5_ORYSJ | LOC_Os08g02230 | 2 | 0.042 | 1.812 |
|  | anionic peroxidase | Q7XSV2_ORYSJ | LOC_Os04g59150 | 11 | 0.005 | 1.491 |
|  | protein disulfide isomerase | A0A0P0XZP3_ORYSJ | LOC_Os11g09280 | 9 | 0.001 | 1.232 |
|  | lactoylglutathione lyase | LGUL_ORYSJ | LOC_Os08g09250 | 11 | 0.000 | 1.193 |
|  | superoxide dismutase | SODC1_ORYSJ | LOC_Os03g22810 | 4 | 0.030 | 1.165 |
|  | superoxide dismutase [Cu-Zn] 4A | SODC2_ORYSJ | LOC_Os07g46990 | 3 | 0.034 | 1.081 |
|  | ferredoxin | FER1_ORYSJ | LOC_Os08g01380 | 3 | 0.038 | 1.054 |
|  | peroxidase 1 | Q5U1S8_ORYSJ | LOC_Os01g22352 | 6 | 0.004 | 0.996 |
|  | peroxiredoxin-2C | PRX2C_ORYSJ | LOC_Os01g48420 | 7 | 0.009 | 0.915 |
|  | ferredoxin-NADP^+^ reductase, embryo isozyme, chloroplastic | FENR3_ORYSJ | LOC_Os07g05400 | 2 | 0.001 | 0.893 |
|  | probable NADPH:quinone oxidoreductase 1 | NQR1_ORYSJ | LOC_Os01g72430 | 3 | 0.002 | 0.891 |
|  | glutathione reductase, cytosolic | GSHRC_ORYSJ | LOC_Os02g56850 | 9 | 0.030 | 0.792 |
|  | Cu/Zn superoxide dismutase (chloroplast) | SODCP_ORYSJ | LOC_Os08g44770 | 7 | 0.015 | 0.718 |
|  | cationic peroxidase SPC4-like | Q5JMS4_ORYSJ | LOC_Os01g73170 | 10 | 0.013 | 0.652 |
|  | thioredoxin H-type | Q6L4X5_ORYSJ | LOC_Os05g43252 | 2 | 0.034 | 0.615 |
| **General metabolic processes** | S-adenosylmethionine synthase 3 | METK3_ORYSJ | LOC_Os01g18860 | 12 | Unique Infested | Unique Infested |
|  | CBS domain-containing protein CBSX1, chloroplastic | Q6YYV0_ORYSJ | LOC_Os09g02710 | 2 | 0.004 | 1.492 |
|  | phosphoserine aminotransferase 2, chloroplastic-like | Q8LMR0_ORYSJ | LOC_Os03g06200 | 5 | 0.008 | 1.325 |
|  | soluble inorganic pyrophosphatase | Q7XTL6_ORYSJ | LOC_Os04g59040 | 5 | 0.006 | 1.189 |
|  | D-3-phosphoglycerate dehydrogenase 1, chloroplastic | Q7XMP6_ORYSJ | LOC_Os04g55720 | 2 | 0.046 | 1.108 |
|  | nucleoside diphosphate kinase 1 | NDK1_ORYSJ | LOC_Os07g30970 | 7 | 0.015 | 0.993 |
|  | nucleoside diphosphate kinase 1 | Q7XC37_ORYSJ | LOC_Os10g41410 | 5 | 0.008 | 0.875 |
|  | probable methylenetetrahydrofolate reductase | MTHR_ORYSJ | LOC_Os03g60090 | 9 | 0.026 | 0.860 |
|  | cysteine synthase | CYSK1_ORYSJ | LOC_Os12g42980 | 15 | 0.003 | 0.779 |
|  | 6-phosphogluconate dehydrogenase, decarboxylating 1 | 6PGD1_ORYSJ | LOC_Os06g02144 | 13 | 0.005 | 0.652 |
|  | adenine phosphoribosyltransferase 1 | Q2QMV8_ORYSJ | LOC_Os12g39860 | 2 | 0.003 | 0.627 |
|  | nucleoside diphosphate kinase 3-like | Q5TKF4_ORYSJ | LOC_Os05g51700 | 3 | 0.001 | 0.613 |
|  | cysteine synthase | Q7XS58_ORYSJ | LOC_Os04g08350 | 2 | 0.009 | 0.610 |
|  | glyceraldehyde-3-phosphate dehydrogenase 1, cytosolic | G3PC1_ORYSJ | LOC_Os08g03290 | 12 | 0.004 | 0.565 |
|  | reactive Intermediate Deaminase A, chloroplastic | Q8H4B9_ORYSJ | LOC_Os07g33240 | 4 | 0.037 | 0.550 |
|  | adenylate kinase 4 | KAD4_ORYSJ | LOC_Os11g20790 | 5 | 0.038 | 0.521 |
|  | 2,3-dimethylmalate lyase-like | Q7XLP7_ORYSJ | LOC_Os04g31700 | 5 | 0.021 | 0.520 |
|  | cysteine synthase | CYSK2_ORYSJ | LOC_Os03g53650 | 6 | 0.003 | 0.513 |
| **Carbohydrate metabolism and energy production** | glucose-1-phosphate adenylyltransferase large subunit, chloroplast precursor, putative, expressed | GLGS1_ORYSJ | LOC_Os09g12660 | 4 | Unique Infested | Unique Infested |
|  | NADP-dependent malic enzyme, chloroplastic | MAOC_ORYSJ | LOC_Os01g09320 | 3 | Unique Infested | Unique Infested |
|  | UTP--glucose-1-phosphate uridylyltransferase | Q6ZGL5_ORYSJ | LOC_Os02g02560 | 5 | Unique Infested | Unique Infested |
|  | probable 6-phosphogluconolactonase 4, chloroplastic | B9G4P3_ORYSJ | LOC_Os09g35970 | 6 | 0.014 | 1.517 |
|  | aspartic proteinase | ASPRX_ORYSJ | LOC_Os05g04630 | 5 | 0.001 | 1.321 |
|  | enolase | Q10P35_ORYSJ | LOC_Os03g14450 | 13 | 0.000 | 0.852 |
|  | 2,3-bisphosphoglycerate-independent phosphoglycerate mutase | Q5QMK7_ORYSJ | LOC_Os01g60190 | 14 | 0.002 | 0.754 |
|  | NAD-dependent epimerase/dehydratase | Q852A3_ORYSJ | LOC_Os03g60740 | 3 | 0.015 | 0.678 |
|  | UDP-glucose pyrophosphorylase | Q93X08_ORYSJ | LOC_Os09g38030 | 16 | 0.024 | 0.637 |
|  | enolase | ENO_ORYSJ | LOC_Os10g08550 | 18 | 0.001 | 0.615 |
|  | phosphoglycerate kinase, cytosolic | Q6H6C7_ORYSJ | LOC_Os02g07260 | 10 | 0.014 | 0.605 |
|  | NADP-dependent malic enzyme isoform X1 | A0A0P0V7M3_ORYSJ | LOC_Os01g52500 | 9 | 0.007 | 0.501 |
| **Protein modification/degradation** | aspartic proteinase nepenthesin-1-like | Q8LNN1_ORYSJ | LOC_Os10g39260 | 5 | 0.000 | 2.213 |
|  | aspartyl protease family protein At5g10770 | Q69QQ2_ORYSJ | LOC_Os09g30414 | 3 | 0.000 | 1.713 |
|  | oryzain alpha chain | ORYA_ORYSJ | LOC_Os04g55650 | 4 | 0.002 | 1.340 |
|  | peptidylprolyl isomerase ROF1-like | Q657L6_ORYSJ | LOC_Os01g38229 | 2 | 0.005 | 1.069 |
|  | probable carboxylesterase 15 | Q8GSJ3_ORYSJ | LOC_Os07g06830 | 4 | 0.010 | 0.754 |
|  | ubiquitin-conjugating enzyme E2 36 | Q8W0I1_ORYSJ | LOC_Os01g48280 | 3 | 0.026 | 0.733 |
|  | putative Calreticulin precursor | CALR_ORYSJ | LOC_Os07g14270 | 4 | 0.004 | 0.719 |
|  | proteasome subunit beta type-2 | PSB2_ORYSJ | LOC_Os03g48930 | 4 | 0.004 | 0.663 |
|  | peptidyl-prolyl cis-trans isomerase | Q6ZH98_ORYSJ | LOC_Os02g02890 | 5 | 0.002 | 0.538 |
|  | proteasome subunit alpha type-5 | PSA5_ORYSJ | LOC_Os11g40140 | 2 | 0.000 | 0.526 |
| **Stress response** | chitinase 8 | CHI8_ORYSJ | LOC_Os10g39680 | 3 | 0.003 | 4.267 |
|  | pathogenesis-related protein 1 | Q6YT69_ORYSJ | LOC_Os07g03730 | 3 | 0.000 | 3.623 |
|  | pathogenesis-related maize seed protein | Q8W084_ORYSJ | LOC_Os01g28500 | 3 | 0.013 | 2.090 |
|  | osmotin-like protein | Q2QND6_ORYSJ | LOC_Os12g38170 | 5 | 0.004 | 1.524 |
|  | ricin B-like lectin R40G2 | 40G2_ORYSJ | LOC_Os07g48490 | 9 | 0.001 | 1.290 |
|  | xylanase inhibitor protein 2-like | Q5WMX0_ORYSJ | LOC_Os05g15770 | 7 | 0.013 | 1.107 |
|  | heat shock cognate 70 kDa protein 2 | Q84TA1_ORYSJ | LOC_Os03g60620 | 11 | 0.015 | 0.999 |
|  | ricin B-like lectin R40G3 | Q9FTY4_ORYSJ | LOC_Os01g01450 | 2 | 0.010 | 0.997 |
|  | chitin elicitor-binding protein-like | CEBIP_ORYSJ | LOC_Os03g04110 | 4 | 0.001 | 0.936 |
|  | stress-response A/B barrel domain-containing protein UP3 | A0A0P0X8Q0_ORYSJ | LOC_Os07g41820 | 2 | 0.001 | 0.926 |
|  | osmotin-like protein | Q2QND8_ORYSJ | LOC_Os12g38150 | 3 | 0.037 | 0.741 |
|  | chalcone--flavonone isomerase | CFI_ORYSJ | LOC_Os03g60509 | 6 | 0.050 | 0.652 |
|  | heat shock cognate 70 kDa protein 2 | Q10NA9_ORYSJ | LOC_Os03g16860 | 14 | 0.050 | 0.623 |
| **Translation** | putative RNase S-like protein precursor | Q69JF3_ORYSJ | LOC_Os09g36700 | 5 | 0.000 | 2.267 |
|  | ribonuclease 3 | Q69JX7_ORYSJ | LOC_Os09g36680 | 11 | 0.001 | 1.669 |
|  | glycine-rich RNA binding protein | A0A0P0W1Y6_ORYSJ | LOC_Os03g46770 | 6 | 0.010 | 1.014 |
|  | glycine-rich RNA binding protein | Q2QLR2_ORYSJ | LOC_Os12g43600 | 5 | 0.004 | 0.857 |
|  | ribosome-recycling factor, chloroplastic | RRFC_ORYSJ | LOC_Os07g38300 | 4 | 0.030 | 0.636 |
|  | elongation factor 1-delta 1 | EF1D1_ORYSJ | LOC_Os07g42300 | 4 | 0.000 | 0.566 |
|  | ADP-ribosylation factor 2 | ARF2_ORYSJ | LOC_Os05g41060 | 10 | 0.027 | 0.564 |
|  | elongation factor 1-delta-like | EF1B_ORYSJ | LOC_Os07g46750 | 7 | 0.040 | 0.530 |
| **Lipid metabolism** | lipoxygenase 2.1, chloroplastic | Q2QNN5_ORYSJ | LOC_Os12g37260 | 16 | 0.001 | 4.707 |
|  | non-specific lipid-transfer protein 1 | Q2QYL0_ORYSJ | LOC_Os12g02330 | 2 | 0.017 | 2.275 |
|  | lipid transfer protein | NLT2B_ORYSJ | LOC_Os12g02310 | 3 | 0.035 | 1.253 |
|  | 3-ketoacyl-CoA thiolase 2, peroxisomal | Q94LR9_ORYSJ | LOC_Os10g31950 | 3 | 0.015 | 1.003 |
|  | probable plastid-lipid-associated protein 6, chloroplastic | Q2R1S1_ORYSJ | LOC_Os11g38260 | 5 | 0.014 | 0.674 |
|  | acyl transferase 9 | Q9FTG9_ORYSJ | LOC_Os01g42880 | 3 | 0.018 | 0.501 |
| **Photosynthesis** | psbP domain-containing protein 4, chloroplastic | Q2QWM6_ORYSJ | LOC_Os12g08830 | 6 | 0.036 | 0.628 |
|  | ruBisCO large subunit-binding protein subunit beta, chloroplastic | Q6ZFJ9_ORYSJ | LOC_Os02g01280 | 5 | 0.014 | 0.544 |
|  | psbP domain-containing protein 1, chloroplastic | Q2QNI4_ORYSJ | LOC_Os12g37710 | 2 | 0.045 | 0.517 |
| **Protease inhibitor** | putative Bowman Birk trypsin inhibitor | A5HEI2_ORYSJ | LOC_Os01g03340 | 4 | 0.026 | 3.248 |
|  | putative Bowman Birk trypsin inhibitor | Q0JR29_ORYSJ | LOC_Os01g03310 | 3 | 0.001 | 3.024 |
| **Hormone-related** | allene oxide cyclase, chloroplastic | AOC_ORYSJ | LOC_Os03g32314 | 3 | 0.002 | 1.260 |
|  | putative Tryptophan synthase beta chain | Q67VM2_ORYSJ | LOC_Os06g42560 | 4 | 0.003 | 1.032 |
| **Amino acid metabolism** | aspartate aminotransferase, cytoplasmic | AATC_ORYSJ | LOC_Os01g55540 | 2 | 0.010 | 0.846 |
|  | peptide methionine sulfoxide reductase A4, chloroplastic | MSRA4_ORYSJ | LOC_Os10g41400 | 4 | 0.006 | 0.811 |
| **Others** | temperature-induced lipocalin-1 | Q6K623_ORYSJ | LOC_Os02g39930 | 4 | 0.000 | 1.911 |
|  | legumin-like protein | Q65XA1_ORYSJ | LOC_Os05g02520 | 6 | 0.003 | 0.798 |
|  | mannose/glucose-specific lectin | Q306J3_ORYSJ | LOC_Os12g14440 | 11 | 0.000 | 4.968 |
|  | transaldolase 2 | Q5JK10_ORYSJ | LOC_Os01g70170 | 12 | 0.003 | 1.273 |
|  | PLAT domain-containing protein 3 | Q6ZGP5_ORYSJ | LOC_Os02g51710 | 4 | 0.009 | 1.261 |
|  | PLAT domain-containing protein 3 | Q7XRE7_ORYSJ | LOC_Os04g38390 | 4 | 0.002 | 1.185 |
|  | putative F8K7.10 protein | Q6AVR6_ORYSJ | LOC_Os03g62370 | 4 | 0.001 | 1.179 |
|  | protein C2-DOMAIN ABA-RELATED 8-like | Q69RN2_ORYSJ | LOC_Os07g31720 | 3 | 0.009 | 0.907 |
|  | luminal-binding protein 2 | BIP1_ORYSJ | LOC_Os02g02410 | 7 | 0.002 | 0.617 |
|  | putative PrMC3 | A0A0P0WTX9_ORYSJ | LOC_Os06g11135 | 4 | 0.026 | 0.613 |
|  | heme-binding protein 2 | Q9LD82_ORYSJ | LOC_Os01g11230 | 6 | 0.010 | 0.609 |
|  | 14-3-3-like protein | 14336_ORYSJ | LOC_Os03g50290 | 12 | 0.010 | 0.533 |
